# Expression bias in retinoic acid responsive genes defines variations in neural differentiation of human pluripotent stem cells

**DOI:** 10.1101/2021.03.17.435870

**Authors:** Suel-Kee Kim, Seungmae Seo, Genevieve Stein-O’Brien, Amritha Jaishankar, Kazuya Ogawa, Nicola Micali, Victor Luria, Amir Karger, Yanhong Wang, Thomas M. Hyde, Joel E. Kleinman, Ty Voss, Elana J. Fertig, Joo-Heon Shin, Roland Bürli, Alan J. Cross, Nicholas J. Brandon, Daniel R. Weinberger, Joshua G. Chenoweth, Daniel J. Hoeppner, Nenad Sestan, Carlo Colantuoni, Ronald D. McKay

**Affiliations:** Lieber Institute for Brain Development, 855 North Wolfe Street, Baltimore, MD 21205, USA; Department of Neuroscience, Kavli Institute for Neuroscience, Program in Cellular Neuroscience, Neurodegeneration and Repair, Child Study Center, Yale School of Medicine, New Haven, CT 06510, USA; Departments of Genetics, of Psychiatry, and of Comparative Medicine, Kavli Institute for Neuroscience, Program in Cellular Neuroscience, Neurodegeneration and Repair, Child Study Center, Yale School of Medicine, New Haven, CT 06510, USA; Department of Systems Biology, Harvard Medical School, Boston, MA 02115, USA; Division of Genetics and Genomics, Boston Children’s Hospital, Harvard Medical School, Boston, MA 02115, USA; IT-Research Computing, Harvard Medical School, Boston, MA 02115, USA; Department of Cell Biology, Johns Hopkins School of Medicine, Baltimore, MD 21205, USA; Department of Neurology, Johns Hopkins School of Medicine, Baltimore, MD 21205, USA; Departments of Oncology, Biomedical Engineering, and Applied Mathematics and Statistics, Johns Hopkins School of Medicine, Baltimore, MD 21205, USA; Department of Psychiatry, Johns Hopkins School of Medicine, Baltimore, MD 21205, USA; Department of Neuroscience, Johns Hopkins School of Medicine, Baltimore, MD 21205, USA; McKusick-Nathans Institute of Genetic Medicine, Johns Hopkins School of Medicine, Baltimore, MD 21205, USA; Division of Preclinical Innovation, Nation Center for Advancing Translational Science, NIH, Bethesda, MD 20892, USA; Astra-Zeneca Neuroscience iMED., 141 Portland Street, Cambridge, MA 01239, USA; Institute for Genome Sciences, University of Maryland School of Medicine, Baltimore, MD 21201, USA

**Keywords:** Pluripotent stem cells, Embryonic development, Retinoic acid signaling, Variation in neural fates, Human variation, Epigenetic control, SOX21, GBX2

## Abstract

Variability between human pluripotent stem cell (hPSC) lines remains a challenge and opportunity in biomedicine. We identified differences in the early lineage emergence across hPSC lines that mapped on the antero-posterior axis of embryonic development. RNA-seq analysis revealed dynamic transcriptomic patterns that defined the emergence of mesendodermal versus neuroectodermal lineages conserved across hPSC lines and cell line-specific transcriptional signatures that were invariant across differentiation. The stable cell line-specific transcriptomic patterns predicted the retinoic acid (RA) response of the cell lines, resulting in distinct bias towards fore-versus hind-brain fates. Replicate hPSC lines and paired adult donor tissue demonstrated that cells from individual humans expressed unique and long-lasting transcriptomic signatures associated with evolutionarily recent genes. In addition to this genetic contribution, we found that replicate lines from a single donor showed divergent brain regional fates linked to distinct chromatin states, indicating that epigenetic mechanisms also contribute to neural fate differences. This variation in lineage bias and its correlation with RA responsive gene expression was also observed in a large collection of hPSC lines. These results define transcriptomic differences in hPSCs that initiate a critical early step specifying anterior or posterior neural fates.

## Introduction

In mammalian embryos, the few hundred pluripotent cells of the epiblast execute spatially coordinated restrictions in cell state, generating distinct tissues (1, 2). Human pluripotent stem cells (hPSCs) represent the epiblast state, acutely poised to diversify into the embryonic germ layers, including endoderm, mesoderm, and ectoderm, before generating all the major organs (3–6). Great attention is currently focused on using hPSCs to generate disease models that promise novel drug and cell therapies (7). Previous studies have defined variation in pluripotent cells by exploring the genomes and transcriptomes of many lines (8–13). However, we lack a detailed understanding of variation in the transitions from pluripotent cells into neural stem cells with distinct brain regional identities, complicating the application of stem cell technologies to neurological and psychiatric disorders. Advances in methods to direct hPSCs to construct human tissues *in vitro* are advancing at an unprecedented rate. These efforts have necessarily focused on developing protocols that generate reproducible cellular output from diverse lines. It is now of great interest to develop assays that allow the exploration of the origins of inherent variation between hPSC lines.

The urgency to address this need is illustrated by recent work reporting developmental differences between hPSC lines in generating regionally specified neural precursors and their possible implications in the etiology of neurodevelopmental disorders (14–21). Here we employ cellular and genomic approaches to define functionally relevant variation in hPSC lines during *in vitro* morphogenesis as they traverse the earliest events in forebrain and hindbrain development. High-resolution decomposition of transcription during hPSC differentiation revealed dynamic transcriptomic changes in lineage emergence that are conserved between lines and stable gene expression signatures that are specific to individual hPSC lines and donors. We show that these transcriptomic signatures are regulated by both genetic and epigenetic mechanisms and demonstrate that they influence cellular trajectories in neural development. The multi-omics data that we have generated and the public data that we use to understand our in vitro observations have been integrated into a gene expression environment using the gEAR/NeMO Analytics framework (22) and can be explored at https://nemoanalytics.org/p?l=Kim2023&g=GBX2.

## Results

### Cell line variation in the emergence of neural fate from pluripotency

Recent studies have defined multiple lineage-competent states that generate distinct embryonic and extraembryonic fates on unconstrained (23–25), surface patterned (26, 27), and microfluidic directed (28) hPSC colony organization. Here, we measured the spatial dynamics of early embryonic cell fate emergence in our unconstrained monolayer culture system by monitoring the expression of key lineage regulators and signaling targets. One day after the passage of dissociated single cells (Day 0, D0), ROCK inhibitor was removed to allow the undifferentiated cells to form epithelial sheets. To induce differentiation, hPSCs were exposed to agonists or antagonists of BMP/TGFβ signaling, known to promote the early steps in gastrulation generating mesendodermal and neural fates, respectively (29, 30). Treatment of cells with BMP4 on day 0 (D0T) induced phosphorylation of SMAD1/5 and coexpression of its targets BRACHYURY (TBXT) and CDX2, which mediate differentiation to early primitive streak and extraembryonic fates (30, 31) across the entire cell population (Supplementary Fig. S1*A*). When BMP4 was treated on day 2 (D2T), the induction of these markers was restricted to the edge of the epithelium (Supplementary Fig. S1*A, B*). However, cells in the core remained competent to respond to BMP4, as indicated by induced phosphorylation of SMAD2/3 (Supplementary Fig. S1*B*). Expression of SOX17 and GATA4, which mediate differentiation to primitive and definitive endoderm (30), suggested that these fates occurred in the core zone (Supplementary Fig. S1*B*). When cells were treated with BMP/TGFβ signaling antagonists Noggin and SB431542 (NSB), which induce neural differentiation (29), the neuroectodermal fate regulators SOX2, SOX21, and OTX2 (32, 33) were expressed in the core of the epithelium while cells on the edge still expressed a high level of pluripotency regulator NANOG (Supplementary Fig. S1*C, D*). Here, we first demonstrate that this 2-dimensional unconstrained system recapitulates aspects of early neuroepithelial lineage emergence in a spatially organized way.

This spatial organization has not been previously tested between hPSC lines in a systematic way. To measure the variation in these self-organizing processes between hPSC lines, the hESC line SA01 was compared with the hiPSC line i04 (34). The two lines showed consistent differences in the formation of core and edge zones defined by larger core zones and greater expression of SOX21 and OTX2 in SA01 and larger edge zones and greater NANOG expression in i04 (Fig. 1*A* and Supplementary Fig. S1*E*). This cell line difference was invariant across doses of neuroectodermal inducers and initial cell plating densities (Supplementary Fig. S1*F, G*) and was not due to differential proliferation rate between the lines (data not shown). When variation in BMP4-induced differentiation was assessed, i04 rapidly induced CDX2, while SA01 showed high TBXT expression (Supplementary Fig. S1*G*). These data demonstrate that functional variation in early fate bias can be defined in this system where dissociated pluripotent cells rapidly self-organize to form developmentally distinct fates.

**Fig. 1.**
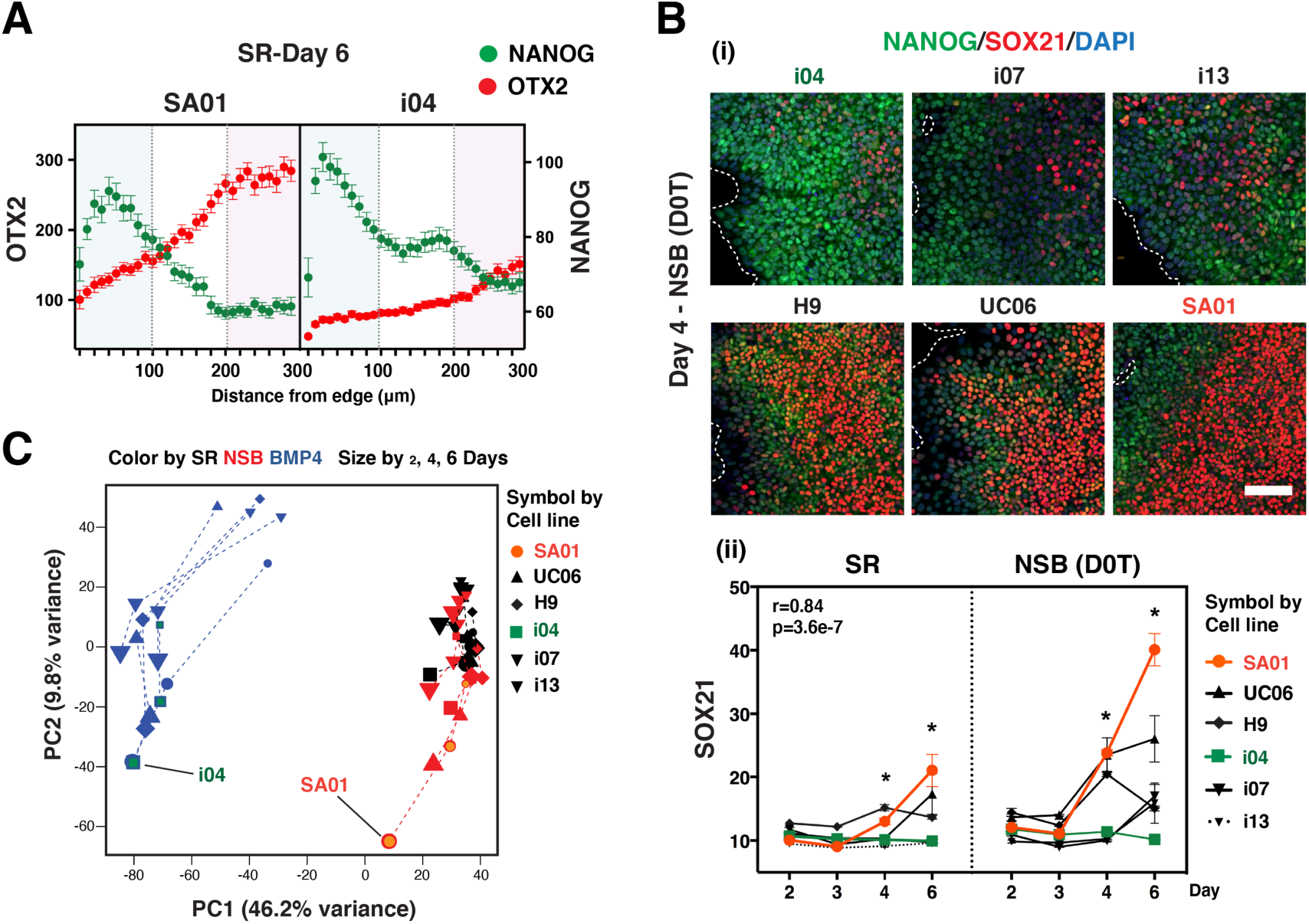
Cell line variation in the emergence of neural fate from pluripotency. (**A**) Spatial expression of NANOG and OTX2 at day 6 in self-renewal (SR) condition in 2 different hPSC lines SA01 and i04. (**B**) Variation in NANOG and SOX21 protein expression across different lines in SR and neuroectoderm (NSB) conditions. (i) Representative images on day 4. Scale bar, 100 μm. (ii) Expression levels in each line plotted across time (n=4, *: Comparison between SA01 and i04: p<0.001). (**C**) PCA of RNA-seq data showing differentiation trajectories.

Variation in the specification of early neural fates between multiple lines was demonstrated by consistent SOX21 protein expression differences between three hESC (H9, UC06, SA01) and three hiPSC lines (i04, i07, i13) in both self-renewal (SR) and neuroectoderm (NSB) conditions (Fig. 1B). RNA sequencing (RNA-seq) was performed at 2, 4, and 6 days of the SR, NSB, and BMP4 conditions for all lines (Supplementary Table S1). In principal component analysis (PCA), PC1 indicated that the major change in gene expression was associated with mesendodermal differentiation and PC2 showed ordered change with differentiation time in all conditions (Fig. 1C). All six lines exhibited similar general trajectories across PC1 and PC2, while cell line differences were also evident. Consistent with the SOX21 expression level, SA01 advanced the furthest along the NSB trajectory (Fig. 1C and Supplementary Fig. S1*H*). This cell line bias in differentiation could also be observed in the PCA of NSB samples alone (Supplementary Fig. S1*I*). To relate the dynamics described by NSB PC1 to expression within SR, we projected the SR data into the transcriptomic space defined by NSB differentiation using projectR (35). The same ranking of cell lines seen in NSB PC1 was present in SR, indicating that elements of gene expression differences defined in differentiation were already present in pluripotency, with SA01 leading the neural trajectory (Supplementary Fig. S1*I*). Similarly, the projection of SR data into BMP4 PC1 also showed a common ranking of cell lines in both SR and differentiation, with i04 leading in this case. These observations suggest that heterogeneity within pluripotency is linked to bias in the early emergence of fates in hPSC lines and that SOX21 expression can be used as an indicator of early neuroectodermal fate specification.

### Decomposing dynamic and stable transcriptomic modules in early differentiation

To more thoroughly analyze the low dimensional transcriptomic change across these cell lines and conditions, we employed the Genome-Wide Coordinated Gene Activity in Pattern Sets (GWCoGAPS) non-negative matrix factorization algorithm (36, 37) to generate a set of 22 patterns that define gene expression dynamics across the samples (Supplementary Fig. S2*A*, GWCoGAPS-I, gene weights in Supplementary Table S2). GWCoGAPS patterns are used in combination to represent the complete expression pattern of each gene, decomposing multiple signals embedded in the expression of individual genes (Supplementary Fig. S2*B*). Through this analysis, two classes of patterns were identified. One class of dynamic patterns defined transcriptomic trajectories that changed over time or condition, and the other cell line-specific patterns defined transcriptomic differences that were stable over time and condition but varied between cell lines (Fig. 2*A*).

**Fig. 2.**
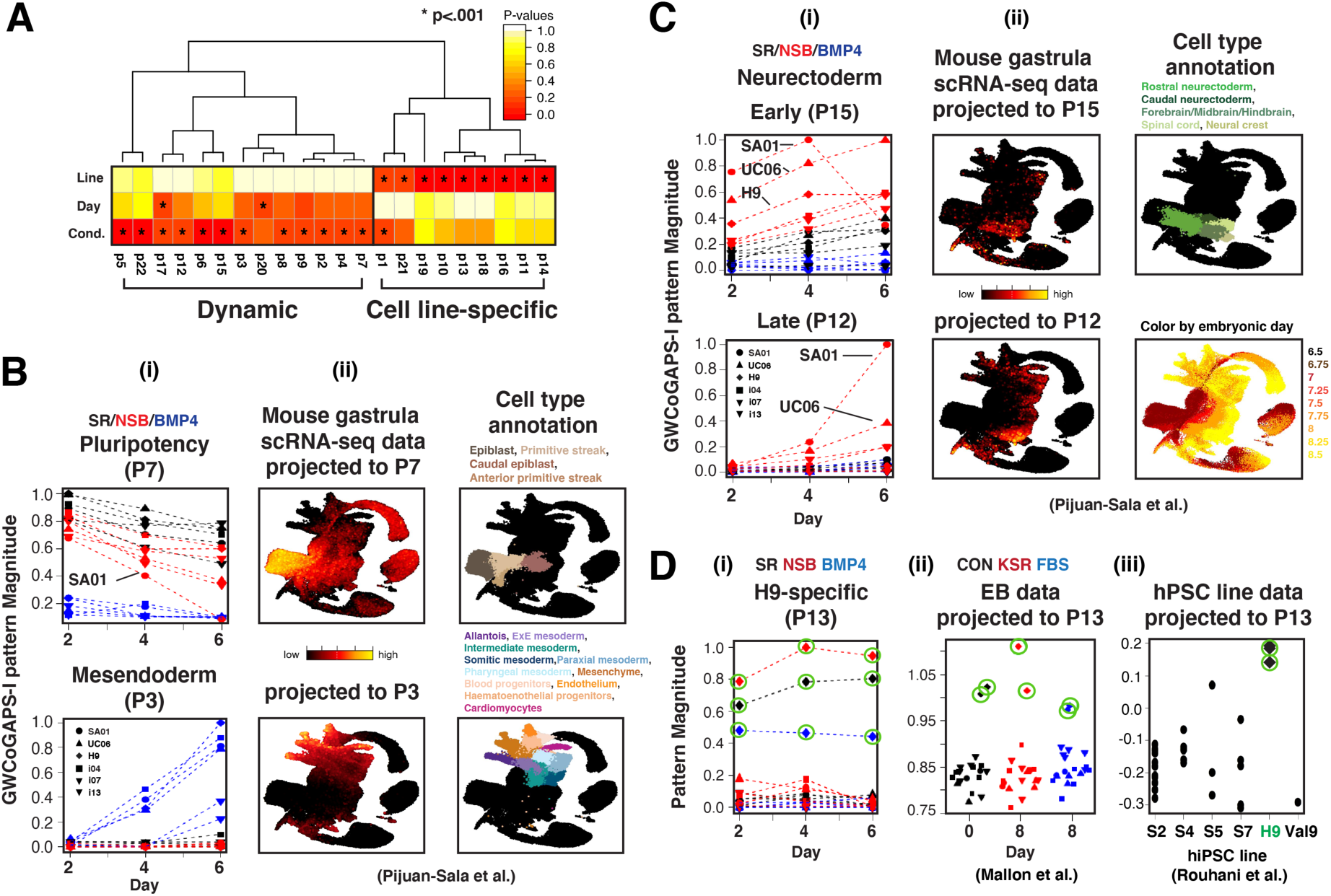
Decomposing dynamic and cell line-specific transcription modules. (**A**) Hierarchical clustering of GWCoGAPS-I patterns and ANOVA p-values for the association of line, time, and condition to these patterns. (**B**) P7 represents loss of pluripotency, and P3 represents induction of mesendoderm regulators. (i) SR pattern P7 and BMP4 pattern P3. (ii) Projections of mouse gastrula scRNA-seq data (38). (**C**) NSB patterns P15 and P12 represent distinct steps in the transition to neural fates. (i) P15 and P12 reveal early and later induction of neuroectoderm differentiation, respectively. (ii) Projection of mouse gastrula data. (**D**) Cell line-specific signatures defined by GWCoGAPS analysis. (i) P13 defines the distinct transcriptomic signature of H9 line from all other lines across time and conditions. (ii) Projection of embryoid body (EB) differentiation data (34) from the same 6 lines (p=3.0e-4). (iii) Projection of multiple hPSC line data (9). H9 samples are circled in green.

Of the 13 dynamic patterns (Fig. 2*A* and Supplementary Fig. S2*A*), three described aspects of pluripotency (P7, P17, and P22), six represented the response to BMP4 (P2, P3, P5, P6, P8, and P9), and three captured different temporal phases of the response to NSB (P4, P15, and P12). Top genes from P7 contained the core pluripotency genes *POU5F1*, *SOX2*, and *NANOG*. Top genes from BMP4 patterns P3 and P9 included early mesendodermal and extraembryonic fate regulators, such as *TBXT, EOMES*, and GATA family members. Top genes from NSB patterns P12 and P15 contained neuroectodermal regulators, such as *HES3*, *WNT1*, *OTX2*, *SOX21*, and *PAX6* (Supplementary Fig. S3*A* and Supplementary Table S3).

To relate these transcriptomic dynamics defined in vitro to gene expression change as lineages emerge during in vivo development, we projected single cell RNA-seq (scRNA-seq) data from the developing mouse gastrula (38) into the GWCoGAPS-I patterns (Fig. 2*B*). Consistent with the designation as a pluripotency module, pattern P7 showed the highest levels in cells of the pluripotent epiblast and decreased in all cell populations undergoing early germ layer commitment. In contrast, the BMP4 pattern P3 was not expressed in pluripotency and increased in later mesodermal lineages, including posterior primitive streak derivatives. Expression of *POU5F1* and *BAMBI*, the top-ranked genes in patterns P7 and P3, respectively, are representative of these expression dynamics in lineage emergence (Supplementary Fig. S3*Bi* and *ii*). Patterns P15 and P12 showed the highest levels in neuroectodermal cells of the mouse embryo (Fig. 2*C*) and parallel expression dynamics of the neural lineage drivers *SOX21* and *PAX6* (Supplementary Fig. S3*Ci* and *ii*). The sequential induction of these early neural expression modules was also found in cortical neuron differentiation data from a different set of hiPSC lines (17) (Supplementary Fig. S3C*iii*).

The GWCoGAPS decomposition also identified cell line-specific expression patterns that were invariant across time and treatment in each cell line (Fig. 2*A*, 2*Di*, and Supplementary Fig. S3*Di*). Projection of microarray data previously generated from embryoid body differentiation of these same hPSC lines (34) showed that these line-specific patterns are stable features found in different culture conditions (Fig. 2*Dii* and Supplementary Fig. S3*Dii*). When RNA-seq data from other research groups using multiple hPSC lines, including the widely studied hESC line H9 (8, 9, 39), were projected into the H9-specific pattern P13, this cell line showed the strongest signal (Fig. 2*Diii* and Supplementary Fig. S3*E*). Projection of DNA methylation data from these same lines showed that cell line-specific gene expression patterns were associated with hypomethylation at promoters overexpressed in the corresponding cell lines (Supplementary Fig. S3*Diii*). These analyses suggest that cell line-specific expression signatures, stable in individual hPSC lines across different conditions, are distinct from dynamic patterns reflecting the transcriptomic change in differentiation.

### SOX21 regulates early forebrain fate by inhibiting mesendoderm and neuromesoderm specification

The high rank of SOX21 in the neuroectoderm initiating pattern P15 (Fig. 2C) and its induction with OTX2 in the NSB condition (Supplementary Fig. S1*D*) suggest its early role in generating forebrain states. Previously, we have shown that SOX21 regulates antero-posterior identity by repressing CDX2 in the adult mouse intestine (32). To better define the role of SOX21 in the early neural specification, three SA01 SOX21-knockout (KO) lines were generated by CRISPR/Cas9 technology (Supplementary Fig. S4*A*). Immunostaining after 3 days of NSB treatment showed increased expression of SOX2 and SOX3 in the epithelial edge zone and increased NANOG in both the edge and core zone of SOX21-KO lines (Fig. 3*A*). These results show the loss of SOX21 disrupts the normal spatial organization regulating the initial emergence of neural fates. RNA-seq data in these KO lines showed that while the overall gene expression pattern conformed to the wild-type (WT) cells, loss of SOX21 delayed neuroectodermal differentiation (Supplementary Fig. S4*B, C*). The expression of the top genes in the pluripotency and neuroectoderm patterns showed that loss of pluripotency gene expression and the transition to neural fates were both diminished in SOX21-KO cells (Supplementary Fig. S4*C*).

**Fig. 3.**
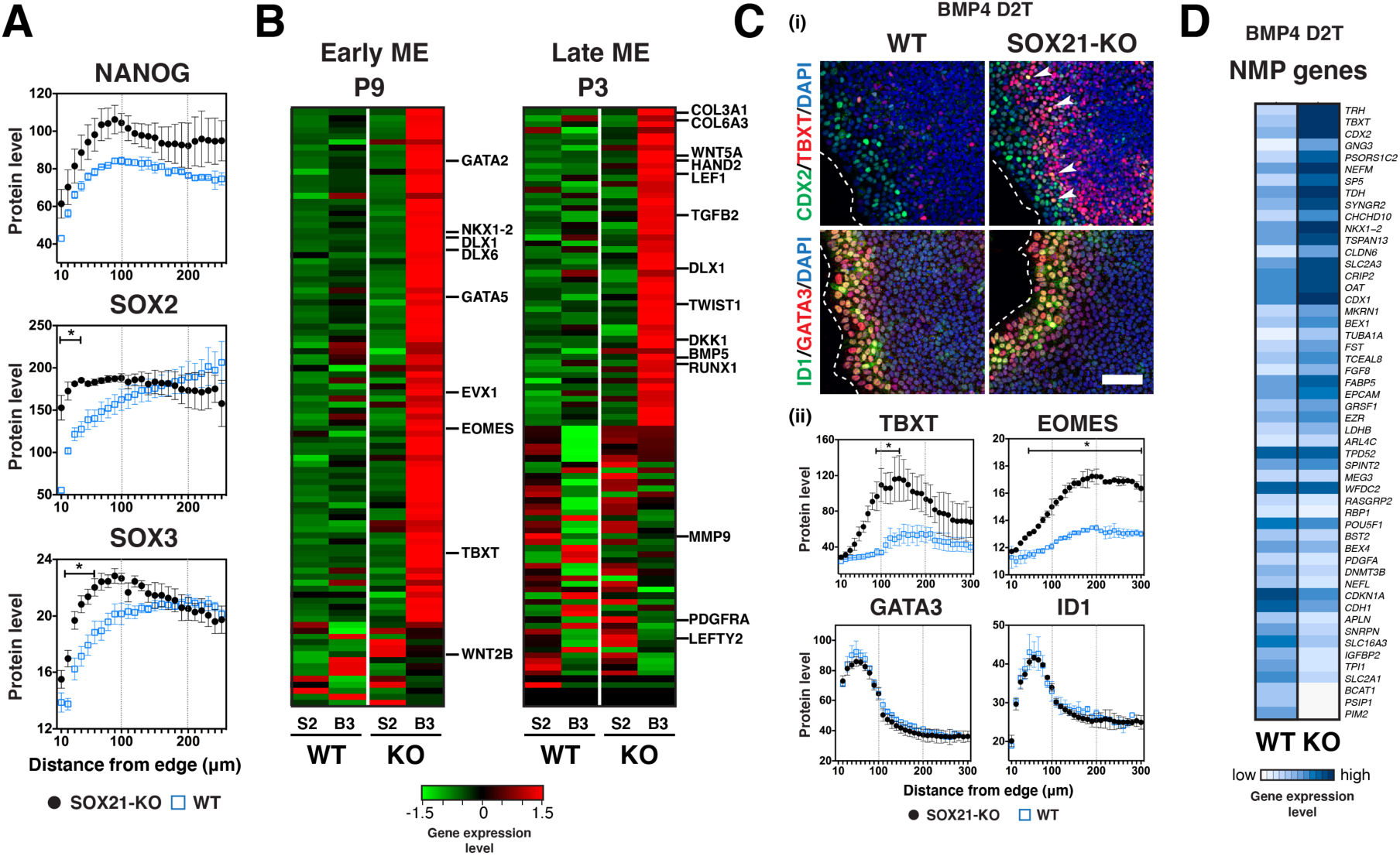
SOX21 mediates early forebrain fate choice. (**A**) Spatial expression of NANOG, SOX2, and SOX3 on day 3 NSB. (**B**) Heatmaps showing the top 100 genes from P9 and P3 on day 2 SR (S2) and at 24 h after BMP4 treatment on day 2 (B3). (**C**) Expression of early primitive streak (TBXT and EOMES) and mesendoderm (CDX2, GATA3, and ID1) regulators at 24 h after BMP4 treatment on day 2 (D2T). (i) Representative images. Arrowheads indicate coexpression of TBXT and CDX2 in SOX21-KO cells. Scale bar, 100 μm. (ii) Spatial expression. *, Comparison between WT (n=3) and SOX21-KO (n=3): p<0.05. (**D**) Heatmap showing the neuromesodermal precursor (NMP) gene expression in WT and SOX21-KO cells at 24 h after BMP4 D2T.

To further test the role of SOX21 in regulating the differential emergence of mesendodermal and neural fates, SOX21-KO cells were treated with BMP4 on day 2 (D2T) when the core zone had already formed in self-renewing cells. Consistent with an early restriction of responsiveness to BMP during the first days of SR (Supplementary Fig. S1*A*), WT cells showed only minimal induction of mesendodermal genes defined by BMP4 patterns P9 and P3 (Fig. 3*B*). In contrast, many BMP4-induced genes were strongly upregulated in SOX21-KO cells. Immunostaining showed no difference between the WT and SOX21-KO lines in the expression of GATA3 and ID1 in the edge zone, while primitive streak regulators TBXT and EOMES (40) were induced strongly and rapidly in the core zone of SOX21-KO cells (Fig. 3*C*). Notably, in the KO cells, CDX2 expression was also extended in the core zone resulting in increased TBXT and CDX2 coexpression. Within the posterior region of embryo, neuromesodermal precursors (NMP) give rise to both neural tissues of spinal cord and trunk mesoderm (41). Consistent with the defined role of CDX2 in specifying NMP (38, 42), the NMP transcriptomic signature was mutually exclusive with SOX21 expression (Supplementary Fig. S4*Dv*). Many genes in the NMP signature showed higher expression in SOX21-KO cells when treated with BMP4 at day 2 (Fig. 3*D*). These results show that SOX21 acts on lineage specification to restrict the expression of genes at the earliest stages of neural fate specification.

This role for SOX21 in early specification of rostral neuroectoderm was also supported by the observation that genes upregulated in SOX21 KO cells in the NSB condition showed higher expression in the epiblast and anterior primitive streak of the developing mouse (Supplementary Fig. S4*Dvi*). Interestingly, the role of SOX21 in restricting caudal identity could be observed by genes upregulated in SOX21-KO cells in BMP4 D2T conditions showing higher expression in caudal epiblast and neural crest. In particular, these data show that the pro-mesodermal and neural crest specifying gene TBXT (43) is among the targets of SOX21 (Fig. 4*D*). Increased *DNMT3A* and decreased *DMNT3B* gene expression in the SOX21-KO lines (https://nemoanalytics.org/p?l=Kim2023&g=DNMT3A) further supports the inhibition of neural crest fate by SOX21 (44, 45). Other laboratories have reported that SOX21 promotes extraembryonic fates at the totipotent mouse 4-cell stage (46), later events in forebrain development using human iPSC model (47), and adult mouse hippocampal neurogenesis (48). These data suggest that SOX21 repeatedly interacts with other class B SOX genes SOX1, 2, and 3 to regulate major morphogenetic decisions. Our data show the role of SOX21 in restricting posterior fates, including NMPs, and promoting anterior neuroectodermal fates as cell emerge from pluripotency.

**Fig. 4.**
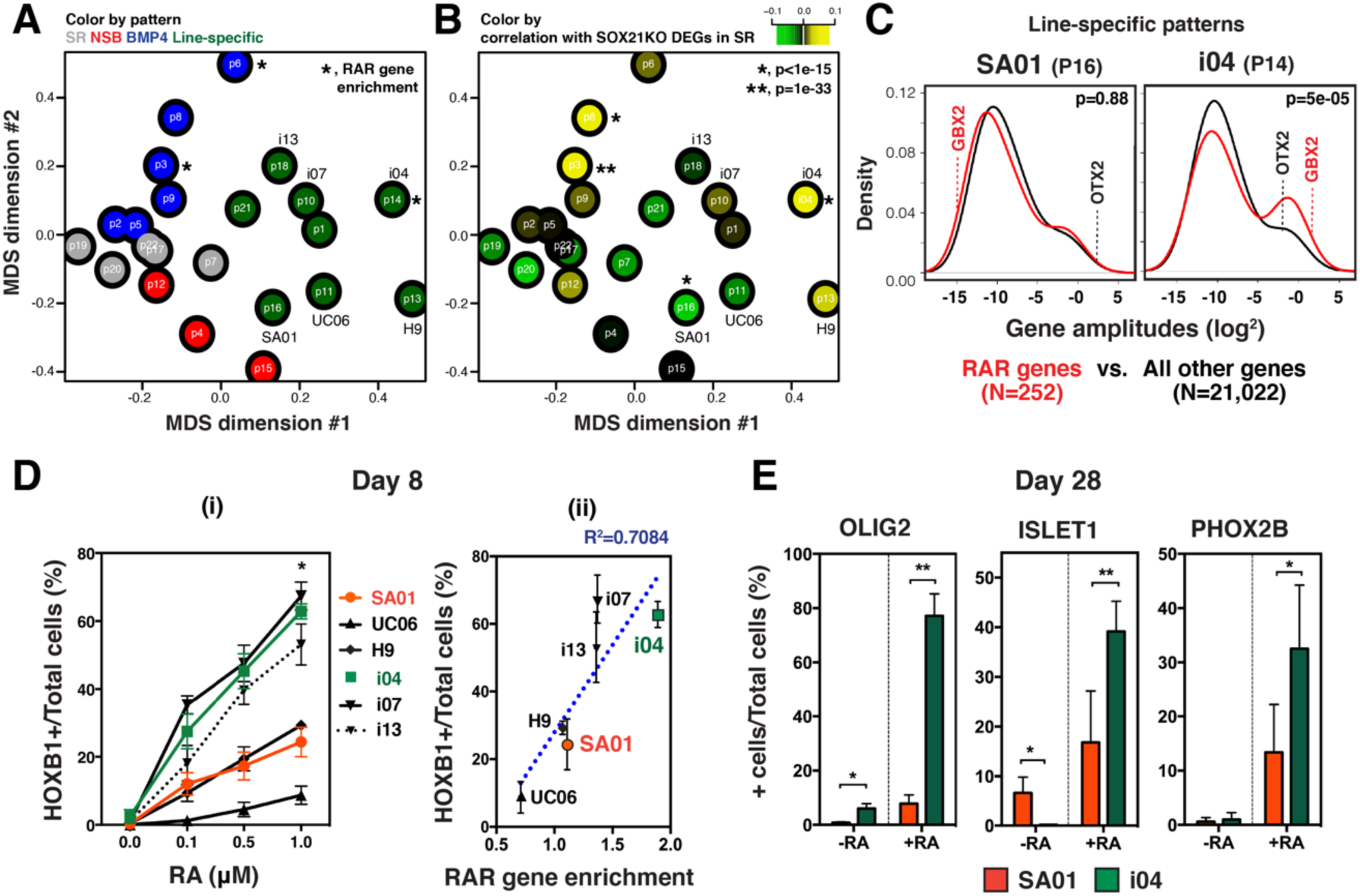
Cell line-specific transcriptomic signatures underlie variation in forebrain versus hindbrain fate bias. (**A**) Multidimensional scaling (MDS) plot of gene amplitudes for the 22 GWCoGAPS-I patterns shows a correlation structure between the patterns. Gray: SR patterns, Red: NSB patterns, Blue: BMP4 patterns, and Green: cell line-specific patterns. *, Patterns with RAR gene enrichment (P3, p=9.7e-08; P6, p=0.014; P14, p=0.008). (**B**) MDS plot of the 22 GWCoGAPS-I patterns colored by the correlation of each pattern’s gene-specific A values with differentially expressed genes (DEGs) in SOX21-KO cells in SR condition. (**C**) Distribution of gene-specific weights of 252 RA responsive genes (49) compared to all 21,022 genes in SA01- and i04-specific transcriptomic signatures. (**D**) Correlation of RA response with RAR gene enrichment scores. (i) Proportion of HOXB1^hi^ cells on day 8. *, Comparison between SA01 and i04 (p<0.05). (ii) Scatter plot of HOXB1^hi^ cell proportions at 1 μM RA and RAR gene enrichment scores in each cell line-specific pattern. Linear fit (R^2^=0.7084). (**E**) Differential production of hindbrain neurons in response to RA in SA01 and i04. Number of OLIG2, ISLET1, and PHOX2B expressing cells on day 28. *, Comparison between SA01 and i04 (p<0.05).

### Cell line-specific transcriptomic changes underlying variation in forebrain versus hindbrain fate bias

To further explore how cell line-specific transcriptomic signatures are related to lineage emergence, we used multidimensional scaling (MDS) to show the correlation between the dynamic and cell line-specific GWCoGAPS patterns (Fig. 4*A*). MDS dimension 1 showed the division of GWCoGAPS analysis into the dynamic patterns and cell line-specific expression signatures. The second MDS dimension separated the three NSB patterns from the six BMP4 patterns, while the SR patterns were positioned between these two groups of lineage-related patterns. Gene expression differences caused by SOX21-KO in pluripotency are positively correlated with gene weights of the two BMP4-induced patterns P3 and P8 (Fig. 4*B*). This is consistent with the SOX21’s role in repressing mesendodermal gene expression. In addition, these SOX21-KO changes were positively correlated with the gene weights in the i04 line-specific signature while being negatively correlated with the SA01 line-specific signature (Fig. 4*B*). These opposing correlations with the distinctly behaving i04 and SA01 lines suggest that line-specific transcriptomic signatures interact with SOX21’s function and the lineage bias of these lines.

To further explore the differential antero-posterior axis patterning in these lines, we interrogated the expression of genes known to be upregulated by retinoic acid (RA) (49). In addition to the known role of RA signaling in posterior mesendodermal, neural, and neuromesodermal development (50–52), the RA responsive (RAR) genes also showed the highest expression in the caudal epiblast and posterior neural fates of the mouse gastrula data (Supplementary Fig. S5*A*). These RAR genes were significantly enriched in two BMP4 patterns and SOX21-KO cells in BMP4 condition (Fig. 4*A* and Supplementary Fig. S5*B*). Notably, the i04 line-specific signature P14 was significantly enriched with RAR genes (Fig. 4*C*) and was positioned among the BMP4 patterns in the MDS dimension 2 (Fig. 4*A*), while the SA01 signature P16 showed no enrichment of RAR genes and appeared amongst the NSB patterns in this dimension. Opposing interactions between OTX2 and GBX2 play an important role in positioning the mid-hindbrain boundary, and GBX2 is expressed anterior hindbrain (53). Interestingly, OTX2 and GBX2 were differentially expressed between SA01 and i04 cells in both SR and neuroectoderm conditions (Supplementary Fig. S5*C, D*). This indicates that RA-driven regional lineage bias may be present at distinct levels in hPSC lines.

The differential enrichment of RAR genes in the cell line-specific expression patterns suggested that these patterns could be used to predict the antero-posterior differentiation behavior of individual cell lines. RA response in neural differentiation was interrogated across multiple cell lines by monitoring the expression of HOXB1, known to drive hindbrain cell fates (54). In a RA dose-response study, lines i04, i07, and i13 produced more HOXB1^+^ cells at day 8 compared to lines SA01, H9, and UC06 (Fig. 4*Di*). RAR gene enrichment in the cell line-specific transcriptomic signatures was strongly correlated with the hindbrain fate potential (Fig. 4*Dii*). Expression of OTX2, GBX2, HOXB1, and HOXB4 in neural precursors at day 8 shows that this differential response in lines i04 and SA01 is consistently observed with different doses of RA (Supplementary Fig. S5*E*). Later differentiation at day 28 showed a coordinated production of multiple hindbrain neuronal fates indicated by expression of OLIG2, ISL1, and PHOX2B (Fig. 4*E*). These data suggest that cell line-specific transcriptomic signatures, especially those associated with RA signaling, regulate the differential emergence of anterior versus posterior neural fates in these hPSC lines.

### Cell line-specific transcription is driven by evolutionarily recent genes under genetic control

To further test the validity of these measures of human cellular variation, a new set of six hiPSC lines were generated from scalp fibroblasts of three donors (2053, 2063, and 2075) whose postmortem brain RNA-seq data we generated previously (55). RNA samples from each donor’s replicate lines were collected at 2, 4, and 6 days under SR and neuroectoderm (LSB) conditions for sequencing. Dynamic differentiation trajectories observed in the original hPSC lines (GWCoGAPS-I) were recapitulated in the analysis of this new RNA-seq data (GWCoGAPS-II, Supplementary Table S4). Extending our observation of line-specific expression signatures, this new analysis identified transcriptomic signatures that were stable across time and conditions for both replicate lines of each donor (Fig. 5*Ai*). Projection of RNA-seq data from the cerebral cortex of 260 individuals (55) into the donor-specific transcriptomic signatures demonstrated that these expression traits were also present in the corresponding donor’s adult postmortem tissue (Fig. 5*Aii*). Projection showed that the donor-specific transcriptomic signatures identified in the pluripotent cells were also highest in the parental fibroblasts of the corresponding donors (Supplementary Fig. S6*A*). These expression data across replicate hiPSC lines, fibroblasts, and mature brain tissue suggest that donor-specific transcriptomic signatures are stable gene expression traits of an individual human’s cells throughout life.

**Fig. 5.**
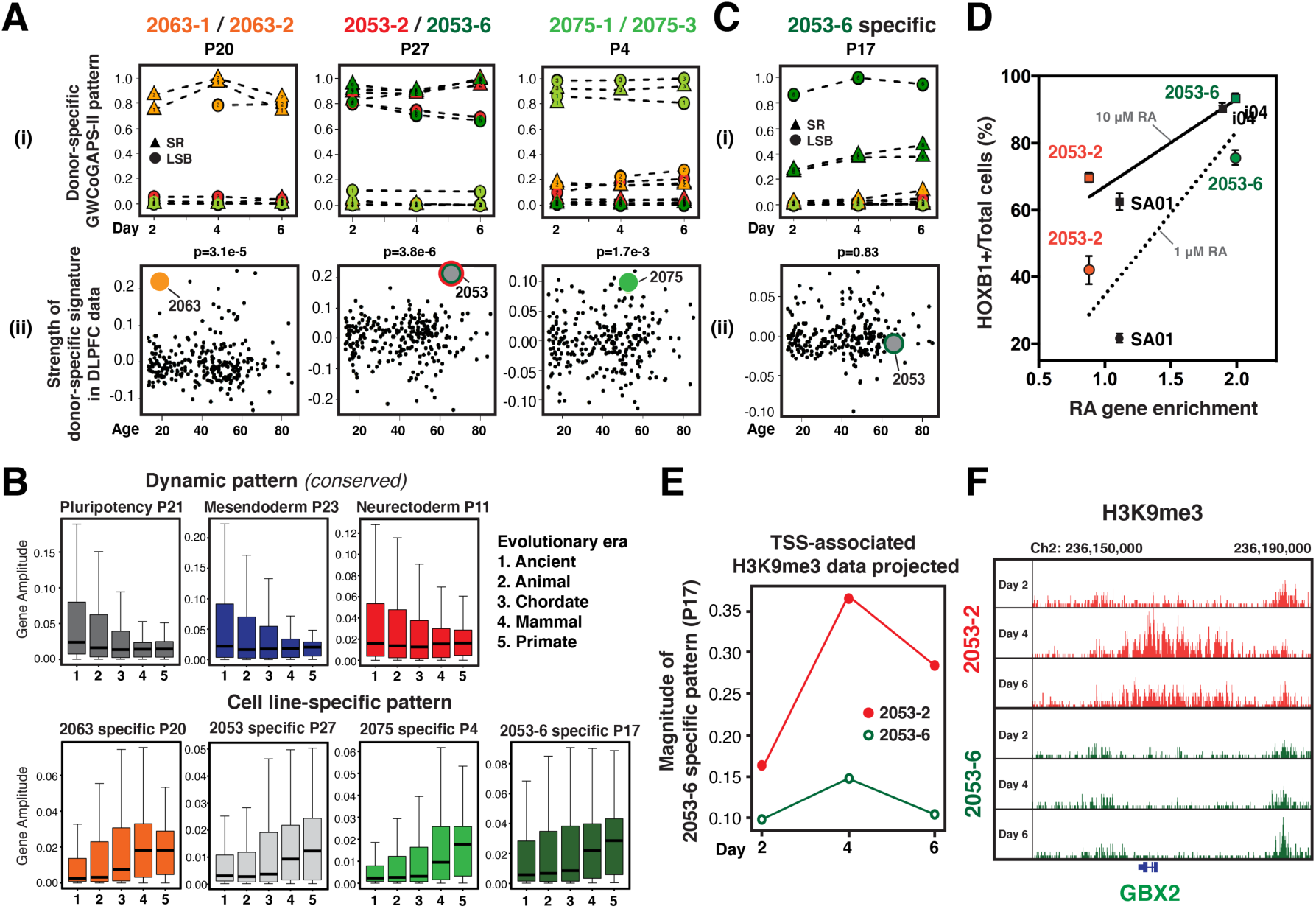
Genetic and epigenetic elements contribute to donor- and line-specific transcriptomic signatures. (**A**) Donor-specific patterns and projections. (i) Patterns representing distinct transcriptomic signatures shared between duplicate lines from the same donor across time and condition. (ii) RNA-seq data of 260 human brain samples projected into each donor-specific pattern. Permutation analysis of these projections showed the significance of the specificity of these donor-specific transcriptomic signatures in mature brain tissue: 2053, p=3.8e-6; 2075, p=1.7e-3; 2063, p=3.1e-5. (**B**) Contribution of genes of different evolutionary eras to GWCoGAPS-II patterns. Conserved dynamic patterns show higher gene amplitudes in ancient genes (comparison on the distribution of era 1 gene amplitudes to era 5 gene amplitudes via Wilcoxon rank sum test: p<1e-16 for all 3 dynamic patterns). Primate-specific genes show higher gene amplitudes in the stable cell line-specific patterns (p<1e-16 for patterns P20 and P27; p<1e-16 for patterns P4 and P7). (**C**) Replicate line-specific variation within a donor. 2053-6 line-specific pattern and projection of brain RNA-seq data. (**D**) Correlation of HOXB1^hi^ cell proportions and RAR gene enrichment scores in each cell line-specific pattern. Linear fit (R^2^=0.75 in 1 μM RA and R^2^=0.87 in 10 μM RA). The HOXB1^hi^ cell proportion in line 2053-2 was correlated with the 2053 donor-specific pattern P27. (**E**) Projection of H3K9me3 ChIP-seq data from lines 2053-2 and 2053-6 in SR into the 2053-6 line-specific transcriptomic pattern. (**F**) H3K9me3 ChIP-seq data in lines 2053-2 and 2053-6 at 2, 4, and 6 days of SR at the GBX2 locus.

The initial Genotype-Tissue Expression (GTEx) consortium data surveyed gene expression and genetic variation in multiple tissues from many donors and identified multi-tissue eQTLs (56). Genes with high amplitudes in our donor-specific signatures were significantly enriched in these eQTLs (p=1e-6 to p=5e-21), further suggesting their stable expression patterns. To assess possible genetic origins of these stable signatures, we compared the strength of the donor-specific signatures in the brain RNA-seq collection with the genetic similarity between donors using single nucleotide polymorphism (SNP) genotypes in these brains (55) (Supplementary Fig. S6*B*). We observed significant (p=4.5e-7 and p=2.5e-8) correlations of genetic similarity with the strength of projected donor-specific transcriptomic signatures, indicating that genetic factors influence donor-specific expression signatures.

This genetic control of variation in gene expression across individual humans may be evolutionarily recent. To test this hypothesis, we estimated the evolutionary ages of genes by phylostratigraphy (57, 58) and examined their distribution across the gene weights of the GWCoGAPS-II patterns. This analysis revealed that dynamic patterns that were consistent among all lines (*conserved*) show stronger weights in evolutionarily ancient genes. In contrast, the dynamic patterns where individual lines showed variation (*divergent*) exhibited a higher contribution of evolutionarily newer genes (Supplementary Fig. S6*C*). Importantly, evolutionarily recent genes showed the highest contribution in stable line-specific transcriptomic signatures (Fig. 5*B*). Similar enrichments were found in the GWCoGAPS-I patterns obtained from the first six hPSC lines (Supplementary Fig. S6*D*). These repeated observations suggest a model where conserved dynamic changes in pluripotency and differentiation are ancient and are part of the widely discussed “Waddington landscape” (59) that constrains cellular differentiation paths. In contrast, the stable transcriptomic patterns relating to individual human variation are newer and influence how the cells of individual hPSC lines follow particular paths within this cellular landscape.

Additionally, estimates of gene dosage sensitivity (60) indicate that genes involved in stable line-specific signatures are less fundamental to survival, consistent with their recent evolutionary origin. We calculated the estimated probability of dosage sensitivity for genes in the top 1% of each of the GWCoGAPS-I patterns and found that gene dosage sensitivity was lower in stable line-specific patterns (average probability of haploinsufficiency, pHaplo, range; 0.32-0.47) than in all other patterns (pHaplo range; 0.53-0.72).

### Cell line-specific transcription can also be driven by early epigenetic mechanisms

In addition to the three donor-specific signatures (Fig. 5*A*), the GWCoGAPS-II patterns revealed a 2053-6 line-specific transcriptomic signature (Fig. 5*C*). Projection of the prefrontal cortex RNA-seq data showed that, unlike donor-specific patterns, this signature was not present in this donor’s brain (Fig. 5*Cii*). Correspondingly, projection of the 2053-6 line-specific signature into the postmortem brain RNA-seq data showed no correlation with genetic distance between donors (Supplementary Fig. S6*B*). These observations suggest that transcriptomic signatures specific to hPSC lines can arise from distinct genetic and epigenetic origins.

To define variation in their neural differentiation trajectories, RNA-seq data from the new lines were projected into the GWCoGAPS-I neuroectodermal patterns P15 and P12 in Fig. 2C. In the neuroectoderm condition, all the new lines showed similar induction of these forebrain patterns, except 2053-6 (Supplementary Fig. S6*E*). Importantly, line 2053-6 showed less SOX21 induction in response to NSB than line 2053-2 (Supplementary Fig. S6*F*), similar to the differences previously observed between lines SA01 and i04 (Fig. 1). Consistent with the regional variation defined in Fig. 4, RAR genes were enriched in the 2053-6 line-specific pattern, and this line, like line i04, generated more HOXB1^hi^ hindbrain cells in response to RA treatment (Fig. 5D and Supplementary Fig. S6*G*). This indicates that the decision to preferentially form forebrain versus hindbrain fates is regulated epigenetically in the 2053-2 and 2053-6 lines.

To rule out the possibility that large-scale somatic or reprogramming-related genomic mutations underlie the observed differentiation bias between the 2053-2 and -6 lines, genome sequencing was performed on the replicate lines and the mature postmortem brain tissue from donors 2053 and 2075. We identified variants gained and lost in each of these genomic DNA samples with respect to the reference genome (Supplementary Fig. S6*H*). For both donors, the majority of copy number variations (CNVs) were shared in all three samples, suggesting minimal genetic alternation during reprogramming. Furthermore, the numbers of CNVs observed uniquely in the 2053 lines did not differ significantly from those in the 2075 lines. These results suggest that the discordant expression traits and lineage bias between lines 2053-2 and -6 are not due to large-scale genome differences.

To find common transcriptomic elements underlying this putative epigenetic lineage bias across the two sets of lines, we compared differences in gene weights between the i04 and SA01 line-specific signatures with differences between the 2053 donor-specific and the 2053-6 line-specific signatures. GBX2 showed high relative expression in lines i04 and 2053-6, which are both biased toward posterior neural fates (Supplementary Fig. S6*I*). This indicates that RAR genes highly expressed in lines i04 and 2053-6, including GBX2, could be responsible for their shared differentiation bias toward posterior fates.

Relevant to possible epigenetic mechanisms that could influence stable gene expression phenotypes, we found that KRAB-ZNF genes were significantly enriched in all line- and donor-specific transcriptomic signatures that we have identified (Supplementary Table S5). KRAB-ZNF genes have been shown to repress the expression of transposable elements during early development and structure lasting H3K9me3-mediated heterochromatin to regulate gene expression in mature tissues (61, 62). These KRAB-ZNF genes are located in gene clusters prone to copy number variation (63). We found that these genes are stably expressed at distinct levels in hPSC lines (Supplementary Fig. S6*J, K*). In addition, consistent with the evolutionary recency of cell line-specific transcriptomic signatures, we observed that human- and primate-specific genes expressed in neural progenitor cells (64), including *ZNF726, ZNF43, ZNF732, ZNF680, ZNF788,* and metallothionein genes *MT1E* and *MT1M*, were expressed in pluripotency and found to be highly ranked in the stable line-specific signatures (Supplementary Table S2 and S4). Notably, differential expression of KRAB-ZNF genes was observed between the lines derived from the same donor. *ZNF676, ZNF208,* and *ZNF729*, previously shown to be highly expressed in human naïve ESCs (65), showed higher expression in 2053-6 lines compared to 2053-2 (Supplementary Fig. S6*J*). Genome sequencing in these lines revealed no detectable CNVs in these or any other differentially expressed KRAB-ZNF genes. These observations raise the possibility that early KRAB-ZNF-driven H3K9me3 and additional epigenetic mechanisms in pluripotency shape lasting transcriptomic phenotypes that influence cell function.

To explore this further, we generated H3K9me3 ChIP-seq data from hiPSC lines 2053-2 and 2053-6 in the SR condition. Projection of these data into the 2053-6 line-specific transcriptomic signature indicated enrichment of this repressive epigenetic mark in line 2053-2 at promoters (+/− 3kb from TSSs) of genes overexpressed in line 2053-6 (Fig. 5*E*). This is consistent with a model in which many specific genes have been derepressed in line 2053-6 via loss of H3K9me3, leading to stable expression phenotypes. GBX2, along with other hindbrain fate genes IRX1/2/4, ZIC1, and OLIG3, were highly represented in the 2053-6 line-specific signature, showed higher H3K9me3 levels in 2053-2 (Fig. 5*F*). In contrast, no consistent H3K9me3 differences were present at the differentially expressed KRAB-ZNF loci (Supplementary Table S6). Also striking was that although HOX genes are not expressed in self-renewal, we observed 35 HOX genes among the top 100 H3K9me3 differences increased in line 2053-2 (Supplementary Table S6). This distinct setting of heterochromatin between the lines in pluripotency, particularly at these canonical regulators of regional order in the metazoan body plan, further supports an early epigenetic origin for their divergent transcriptomic phenotypes and resulting lineage bias.

### Early developmental bias in RA signaling defines hPSC variation in the wider human population

To further explore genetic and epigenetic differences in lineage bias in a larger human population, we performed PCA on RNA-seq data from 317 undifferentiated hiPSC lines from 101 donors generated by the NextGen Consortium (13). As recognized by Carcamo-Orive et al. (13), 91% of the variance in PC1 is attributable to differences across donors, indicating that this dominant feature of human transcriptomic variation within pluripotency is under genetic control (Fig. 6*A*). Importantly, PC1 was highly correlated with RAR gene expression (r=0.79, p=8.9e-70), suggesting that the differences in RA response we have observed in a small number of lines generally apply (Fig. 6*A* and Supplementary Table S7). In addition, similar to the case of lines 2053-2 and 2053-6, a small number of individual donor’s iPSC lines occupied significantly divergent PC1 positions and RAR gene expression levels (Fig. 6*A*, an example in blue circles), confirming that epigenetic mechanisms can also influence this transcriptomic variation.

**Fig. 6.**
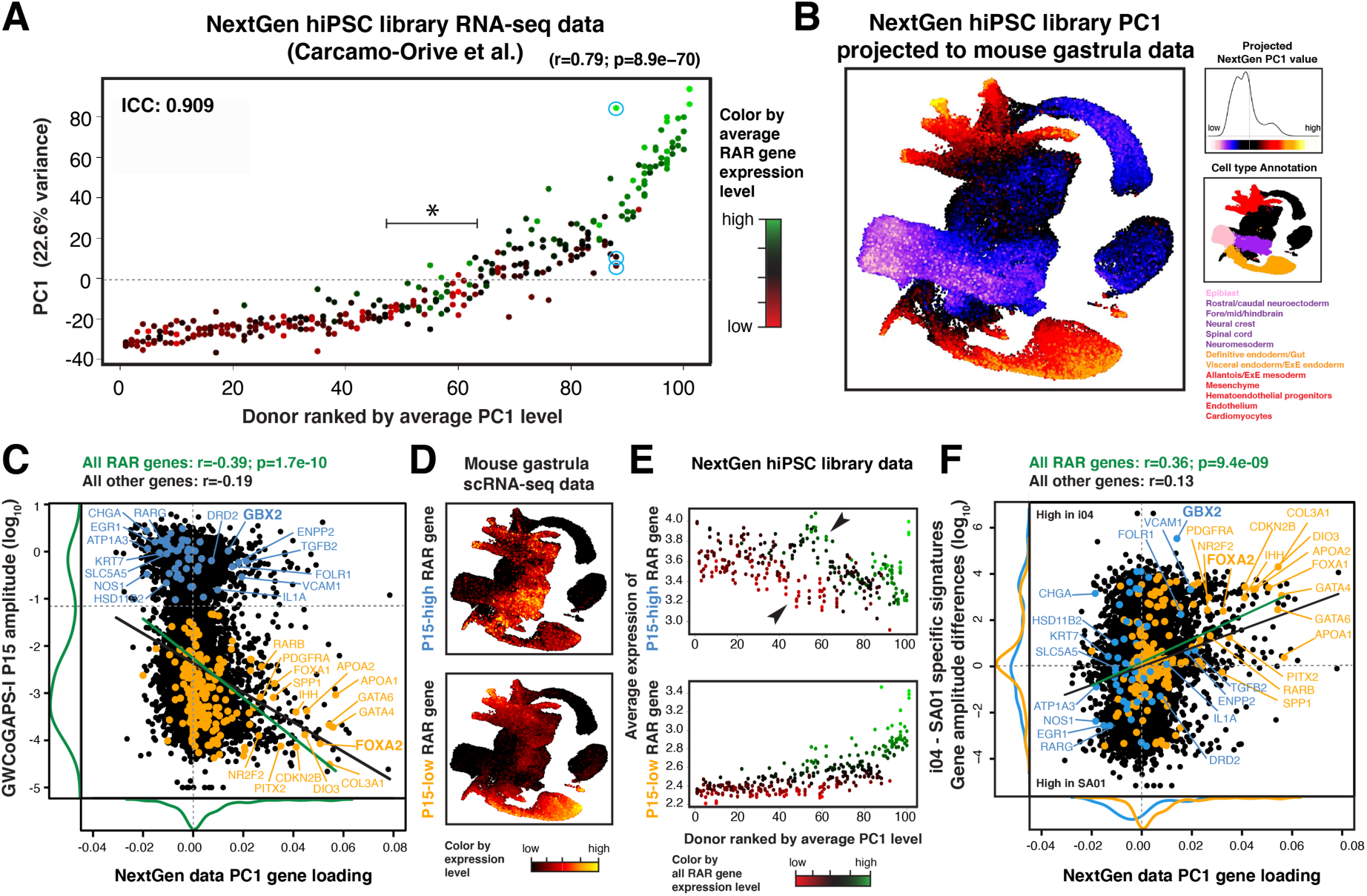
Early developmental bias in the expression of retinoic acid responsive (RAR) genes define hPSC variation in the wider human population. (**A**) PC1 of the NextGen Consortium hiPSC library (317 lines from 101 donors) RNA-seq data (13). Each line is colored by its average RAR gene expression level. Donors are ranked along the X-axis by the average PC1 level of all lines derived from that donor. All lines from one donor are shown vertically along the same X-axis position. Pearson’s R and p-values indicate the correlation of PC1 with mean RAR gene expression. Blue circles indicate lines from one donor, which show a large variance in PC1 and RAR gene expression levels. The PC1 value of zero intersects with an inflection point in the variation across donors. ICC=intraclass correlation coefficient. (**B**) Projection of NextGen PC1 onto the mouse gastrula scRNA-seq data (38), mapping transcriptomic variation in hPSC lines onto in vivo lineage emergence. (**C**) Scatter plot comparing PC1 gene loadings with gene amplitudes of GWCoGAPS-I neuroectoderm pattern P15 (Fig. 2*C*) colored by RAR genes with high (in blue) or low (in yellow) gene amplitudes in P15. (**D**) UMAP plots of mouse gastrula scRNA-seq data colored by the expression level of RAR genes with high or low gene amplitudes in P15. (**E**) Average expression level of RAR genes with high or low P15 gene amplitudes in NextGen RNA-seq data. (**F**) Scatter plot comparing NextGen PC1 gene loadings with differences in gene amplitudes between the i04 and SA01 line-specific patterns colored by RAR genes with high or low gene amplitudes in P15.

Projection of the NextGen PC1 onto the landscape of early mouse development reveals that this variation in hPSC lines contains strong lineage bias (Fig. 6*B*). High PC1 represents both mesodermal (*COL3A1, PRRX1, ANXA1, INHBA*) and endodermal (*RARB, APOA2, CER1, GATA4, GATA3, GATA6, SOX17, EOMES*) identities (Fig. 6*B*, in red to yellow, and Supplementary Table S8). Interestingly, RAR genes showing the highest PC1 loadings included genes shown to be highly expressed in self-renewing cells at the edge of pluripotent colonies (23) and in a recently defined pluripotent founder cell that shares transcriptomic elements with the primitive endoderm (25). In contrast, low PC1 maps to the pluripotent epiblast and emerging neural lineages with high expression of known regulators of anterior neural fates such as *SOX3, LHX2, PAX6, EMX2, FEZF2,* and *RAX* (Fig. 6*B*, in blue to pink, and Supplementary Table S8).

In addition to the high RAR gene expression in hPSC lines with mesendodermal bias in high PC1, a cluster of lines at intermediate PC1 levels also show high RAR gene expression (asterisk in Fig. 6*A*). Projections of GWCoGAPS decomposition of this NextGen data (GWCoGAPS-III, Supplementary Table S8) into the mouse gastrula and our original six hPSC line data indicate that this cluster of cell lines has the early neural bias (Supplementary Fig. S7*Aii*). To further dissect specific RAR genes associated with this neural bias, we examined RAR genes in the NextGen PC1 loadings and the early neuroectoderm pattern P15 of GWCoGAPS-I (Fig. 6*C*). A strong negative correlation between the PC1 loadings and the P15 weights further confirmed that the low NextGen PC1 genes are related to neural lineages. The bimodal distribution of genes in the P15 weights defined two groups of RAR genes (Fig. 6*C*). Average expression of these two RAR gene subgroups in the mouse gastrula revealed neural (P15-high) or mesendodermal (P15-low) signatures (Fig. 6*D*). Interestingly, the average expression of P15-high RAR genes further distinguished two groups of NextGen hiPSC lines within this neural bias (Fig. 6*E*, arrowheads in upper panel). The higher expression of the P15-high RAR genes in the mouse caudal epiblast, caudal neuroectoderm, and spinal cord suggests posterior neural bias in the group of hiPSC lines showing the higher expression of these genes (Supplementary Fig. S7*B*). The analysis of differentially expressed genes between these two groups of hiPSC lines with neural bias revealed differential expression of *GBX2* (Supplementary Fig. S7*C* and Supplementary Table S9) consistent with their posterior neural bias. On the other hand, the average expression of the P15-low RAR genes distinguished hiPSC lines with mesendodermal and primitive endodermal bias, confirmed by *FOXA2* expression (Fig. 6*E*, lower panel and Supplementary Fig. S7*B* and *C*). These results show that the transcriptomic heterogeneity in hPSC lines can be mapped along the developmental axes revealed by the two groups of RAR genes we defined here.

Similarly, to test if cell line-specific signatures also showed a correlation with wider hPSC variation, NextGen PC1 gene loadings were compared to the differences in the stable transcription signatures of the divergently biased lines i04 and SA01 (Fig. 6*F*). We observed a positive correlation when considering all genes (r=0.13) and a stronger correlation within RAR genes (r=0.36, p=9.4e-09). Consistent with a model in which hPSC lines exhibit predictable lineage bias linked to stable transcriptomic traits (Fig. 4), the mesendodermal P15-low RAR genes with high NextGen PC1 loadings show higher levels in the i04-specific signature compared to the SA01-specific signature (Fig. 6*F*, genes in yellow). Our study shows that variation in the lineage bias of hPSC lines can be defined by both dynamic patterns and stable line-specific signatures accessible by the approaches we explore here.

## Discussion

It has been a long-standing interest in stem cell research to predict the differentiation capacity of individual hPSC lines. A growing body of research is revealing the genetic origins of heterogeneity in transcription and differentiation potential of hPSCs (8–14, 39, 66–72). In our work, we define dynamic lineage-driving transcriptomic modules, which are influenced by stable line-specific gene expression traits that hold pluripotent cells in distinct lineage-biased states. The bias we observe in the emergence of the neural lineage from pluripotency in a small number of hPSC lines is also seen in the primary dimension of transcriptomic variation across hundreds of hiPSC lines prior to differentiation. The low variance within donors in the public lines along with our observations in adult tissue suggests that major aspects of this transcriptomic variation in pluripotency are long-lasting and under genetic control.

In addition, our observation of bias toward forebrain versus hindbrain neural fates in replicate lines from the same donor suggests that epigenetic mechanisms also contribute significantly to variation in early neural fate choice. Our data support a model where alternate epigenetically predisposed states within pluripotency exist prior to the high-level transcriptomic output that implements these early regional neural fate choices. This mechanism parallels recent findings in the mouse gastrula, demonstrating epigenetic structuring before transcriptomically defined cell identity (73, 74).

The transcriptomic architecture of self-organization that we define here allows mapping of regional embryonic signatures within hPSCs preceding the full expression of classical organizer cell types. We have previously reported a regional neural fate bias spanning the dorso-ventral telencephalic axis, which was also divergent in hiPSC lines from a single donor (15). We showed that this lineage bias in neural stem cells resulted from the differential emergence of transcriptomic signatures that led to the formation of the dorso-caudal organizer, the cortical hem, or the antero-ventral domain identity. This dorso-ventral specification in neural development has been repeatedly linked to risk for neuropsychiatric disease (16, 21, 75–77). Recent scRNA-seq data in cerebral organoids has demonstrated striking bias in this and other regional neural fate choices across human donors (18, 78). Variation in dorso-ventral specification regulated by differential WNT signaling has also been observed across many hiPSC lines from numerous donors (14). In this study, we define stable cell line-specific expression traits that affect RA signaling to regulate lineage bias along the antero-posterior axis in the transition from pluripotency to neural fates. Dysregulation of RA signaling has been associated with the risk for schizophrenia and autism (51, 79).

In contrast to conserved dynamic expression patterns, we observe that patterns which differ systematically across hPSC lines are also enriched in evolutionarily recent genes. Our results showing KRAB-ZNF gene enrichment and H3K9me3-mediated regulation in cell line-specific signatures suggest possible mechanisms controlling these stable expression phenotypes. Also regulated by H3K9me3 mechanisms, evolutionarily recent transposable elements and their corresponding KRAB-ZNF repressor genes have been directly implicated in human zygotic genome activation and preimplantation states specifying gene expression at later stages of development (65, 80–83)

The genetics of neurodevelopmental disorders (20, 21, 84–86) and human brain evolution (18, 87–89) are being integrated into cerebral organoid models that parallel in utero and early postnatal brain development (90, 91). As PSC-derived models of early human and mouse development become increasingly sophisticated (92–95), approaches to exploring inherent variation across synthetic embryos from diverse PSC lines will become critical. Interest in genetic variation in the early embryo has been substantially enhanced by the observation that there is a selection for euploidy in pluripotency (96). The developmental perspectives we put forward here can inform molecular phenotyping and assay development in these emerging in vitro models.

Recent progress in selecting specific hPSC lines for stem cell therapeutics (70, 71, 97, 98) stresses the continuing value of further defining the genetic and epigenetic control of human cellular variation as hPSCs adopt neural fates. Novel methods that define single-cell genome mosaicism are being combined with lineage tracing (99–104) to provide a new understanding of the importance of cellular selection in neural development (19, 105). New findings indicate the central impact of cellular states within pluripotency on differentiation capacity (106). Our observations raise the question of how many dimensions of hPSC variation can be mapped across in vivo development and how they are mechanistically and functionally interrelated to give rise to diversity in lineage emergence. The functional elements we have defined in pluripotency guide the interrogation of large collections of hiPSC lines from many genetically diverse donors. This work opens the compelling prospect of defining how variation in these early cellular states influences human brain development to modify complex traits and disease risk.

## Materials and Methods

### hPSC culture and differentiation

The hPSCs were dissociated to single cells with accutase (A11105, Life Technologies), plated at a density of 1 X 10^5^ cells/cm^2^ in a Matrigel (354277, BD)-coated plate, and cultured with mTeSR1 (05850, Stem Cell Technology). Cells were plated in medium containing 5 μM Y27632, ROCK inhibitor (Y0503, Sigma-Aldrich) to increase cell survival upon dissociation. ROCK inhibitor was removed from the medium at 24 hours after plating, and cells were cultured for another four days before passaging. For differentiation, cells were plated at a density of 18 X 10^3^ cells/cm^2^ in a Matrigel-coated plate. Noggin (500 ng/ml, 719-NG, R&D Systems) and SB431542 (2 μM, S4317, Sigma-Aldrich) were added to mTeSR1 medium for neuroectodermal differentiation, while BMP4 (100 ng/ml, 314-BP, R&D Systems) was added for mesendodermal differentiation upon ROCK inhibitor removal (Day 0) and cultured for 6 days. For posterior neural differentiation, cells were cultured with Aggrewell medium (05893, Stem Cell Technology) for 2 days after ROCK inhibitor removal and then cultured with N2B27 medium supplemented with LDN193189 (100 nM, 04-0074, Stemgent) and SB431542 (2 μM, S4317, Sigma-Aldrich) for another 4 days. Retinoic acid (R2625, Sigma-Aldrich) was added on day 4 to induce posterior neuroectoderm differentiation. For further differentiation to hindbrain-derived neurons, cells were cultured with neurobasal medium (21103-049, Life Technologies) supplemented with bovine Insulin (25 μg/ml, I6634, Sigma-Aldrich), B27 (17504-044, Life Technologies), human BDNF (10 ng/ml, 248-BD, R&D Systems) and human NT-3 (10 ng/ml, 267-N3, R&D Systems) for another 20 days.

### Generation of hiPSC lines

The hiPSC lines i04, i07, and i13 (NIH-i4, NIH-i7, NIH-i13) have been reported previously (34). The hiPSC lines reprogrammed with synthetic mRNAs were generated using an mRNA reprogramming kit (00-0071, Stemgent) and microRNA Booster kit (00-0073, Stemgent) with modifications. Human fibroblasts (Donor 2075, 2053, and 2063) were seeded at 5 X 10^3^ cells/cm^2^ in a Matrigel-coated plate and cultured with DMEM medium supplemented with 10% FBS (Life Technologies) and 2 mM L-glutamine. After 24 hours (day 1), the medium was changed to Pluriton human NUFF-conditioned media with 300 ng/ml B18R protein. On days 1 and 5, the microRNA booster kit was used with the StemFect RNA transfection reagent kit (00-0069, Stemgent) to enhance reprogramming. On days 2-12, the OSKML RNAs were transfected. The mRNA reprogramming process was performed at 37°C in a 5% O_2_ and CO_2_ incubator.

### Generation of CRISPR/Cas9 mediated SOX21-KO hESC line

The SOX21-KO hESC lines were generated by CRISPR/Cas9 mediated genome deletion system. SOX21-specific gRNAs were designed using the CRISPR Design Tool, Optimized CRISPR Design - MIT for Sox21NHEJ4 (http://crispr.mit.edu/) and CHOPCHOP for Sox21NHEJ5 (https://chopchop.rc.fas.harvard.edu/). The oligonucleotides (CACCGCGGGCTCAGCGGCGCAAGA –top for Sox21NHEJ4; AAACTCTTGCGCCGCTGAGCCCGC –bottom for Sox21NHEJ4; CACCGGGTGTGGTCGCGGGCTCAG –top for Sox21NHEJ5; AAACCTGAGCCCGCGACCACACCC –bottom for Sox21NHEJ5) were cloned into pSpCas9(BB)-2A-Puro (px459; Addgene) and designated the plasmid as pX459-Sox21NHEJ4 and pX459-Sox21NHEJ5. All oligonucleotides were synthesized by Integrated DNA Technologies. SA01 hESCs were transfected with 2.5 μg pX459-Sox21NHEJ4 plasmid or pX459-Sox21NHEJ5 using DNA-In Stem (MTI-Global stem, gifted from Dr. Jessee). Transfected cells were dissociated and plated into a 10 cm culture dish. After 48 hours of 0.5 μg/ml puromycin selection, hESC colonies were maintained for 10 days. Individual colonies were isolated and clonally expanded. Genomic DNA was isolated from each clonal line using Wizard Genomic DNA Purification Kit (Promega). The genomic region surrounding the CRISPR target site for SOX21 was amplified by PCR (KOD Xtreme Hot Start DNA Polymerase; EMD Millipore), and products were treated with SURVEYOR nuclease (SURVEYOR Mutation Detection Kit, Transgenomic) to detect CRISPR/Cas9-induced indel mutations. The PCR products were cloned into pGEM®-T Easy Vector (Promega) and sequenced to confirm the genotypes. Knockout validation of SOX21 protein was performed by immunostaining.

### Immunofluorescence

Cells were fixed with 4% paraformaldehyde for 10 min and permeabilized for 40 min using 0.1% Triton X-100 (Sigma-Aldrich) in PBS. Subsequently, cells were blocked with 10% donkey serum (Sigma-Aldrich) and incubated with primary antibodies overnight. Following primary antibodies and dilutions were used: BRACHYURY (AF2085, R&D, 1:500), CDX2 (AM392, Biogenex), EOMES (ab23345, Abcam, 1:400), GATA3 (MAB6330, R&D, 1:200), GATA4 (AF2606, R&D, 1:400), GBX2 (AF4638, R&D, 1:200), HOXB1 (AF6318, R&D, 1:200), HOXB4 (ab133621, Abcam, 1:400), HOXB9 (ab66765, Abcam, 1:400), ID1 (AF4377, R&D, 1:200), ISLET1 (AF1837, R&D, 1:200), NANOG (AF1997, R&D, 1:200; Reprocell 1:200), OCT4A (MAB17591, R&D, 1:200), OLIG2 (AF2418, R&D, 1:200), OTX2 (AF1979, R&D, 1:200), PAX6 (PRB-278P, BioLegend, 1:500; AF8150, R&D, 1:200), PHOX2B (AF4940, R&D, 1:200), p-SMAD1/5 (9516, Cell Signaling Technology, 1:200), p-SMAD2/3 (8828, Cell Signaling Technology, 1:200), and TUJ1 (PRB-435P, BioLegend, 1:1000), SOX1 (AF3369, R&D, 1:400), SOX17 (AF1924, R&D, 1:500), SOX2 (AF2018, MAB2018, R&D, 1:200), SOX21 (AF3538, R&D, 1:200), SOX3 (GT15119, Neuromics, 1:200), and TUJ1 (MAB1195, R&D, 1:400). Secondary antibody incubation was performed with Alexa flour conjugated antibodies at dilution of 1:400 (Life Technologies). For direct immunostaining, primary antibodies were conjugated using Alexa fluor monoclonal antibody labeling kits (A20181, A20184, A20186, Life Technologies). Nuclei were counterstained with DAPI (Life Technologies).

### High-content spatial analysis

Images were acquired with the Operetta (Perkin Elmer), analyzed in batch mode with custom building blocks on a Columbus server (Perkin Elmer), and visualized with Spotfire (Perkin Elmer). Spatial analysis of hPSC epithelium (‘distance from the edge’ measurement) was achieved using a custom Acapella script (Perkin Elmer) run in Columbus with the following commands; 1) stitch a montage from 3×3 user-defined contiguous overlapping fields captured with the 20x objective, 2) segment and binarize DAPI signal from individual nuclei to create nuclear objects, 3) segment and binarize DAPI signal from the cytoplasm surrounding each nucleus to create cytoplasmic objects. hPSCs show strong blue fluorescence arise from sequestration of retinyl esters in cytoplasmic lipid bodies (107), 4) dilate nuclear objects to eliminate gaps between neighboring objects, 5) create super objects by filling holes containing less than 30 pixels, 6) segment super objects, 7) create a perimeter line at the edge of each super object, 8) calculate the minimum distance between the centroid of each nucleus and the closest super object perimeter, 9) report fluorescence signal from nucleus and cytoplasm for each object. For each cell, this script reports nuclear and cytoplasmic signals for all channels and a single measure of minimum distance to the closest perimeter of the epithelium. Using data visualization in Spotfire, median fluorescence signals from all cells within 10 μm were plotted corresponding to the distance from an edge of epithelium.

### RNA-seq library preparation

Total RNA was extracted using the mirVana kit (Ambion) according to the manufacturer’s protocol. RNA quality control was performed using the Agilent 2100 Bioanalyzer System. RNA-seq libraries were constructed using Illumina mRNA sequencing sample Prep Kit (for Poly-A libraries) or TruSeq Stranded Total RNA RiboZero sample Prep Kit (for strand-specific libraries) following the manufacturer’s protocol. Briefly, poly-A containing mRNA molecules were purified or ribosomal RNAs were removed using RiboZero beads from ~ 800 ng DNase-treated total RNA. Following purification, the resulting RNA was fragmented into small pieces using divalent cations under elevated temperature at 94°C for 2 min. The range of the fragment length was 130-290 bp, with a median length of 185 bp. Reverse transcriptase and random primers were used to copy the cleaved RNA fragments into first-strand cDNA. The second-strand cDNA was synthesized using DNA Polymerase I and RNase H. These cDNA fragments went through an end repair process using T4 DNA polymerase, T4 PNK, and Klenow DNA polymerase, the addition of a single ‘A’ base using Klenow exo (3’ to 5’ exo minus) and the ligation of Illumina PE adapters using T4 DNA Ligase. An index was inserted into Illumina adapters so that multiple samples could be sequenced in one lane of an 8-lane flow cell if necessary. The concentration of RNA was measured by Qubit (Life Technologies). The quality of the RNA-seq library was measured by LabChipGX (Caliper) using HT DNA 1K/12K/HiSens Labchip. The final cDNA libraries were sequenced using HiSeq 2000 (for Poly-A library preparation) or HiSeq 3000 (for RiboZero library preparation) for high-throughput DNA sequencing.

### RNA-seq data processing

After the sequencing, the Illumina Real Time Analysis (RTA) module was used to perform image analysis and base calling, and the BCL Converter (CASAVA v1.8.2) was used to generate FASTQ files, which contain the sequence reads. The sequencing depth was over 80 million (40 million paired-end) mappable sequencing reads (Supplementary Table S1). Read-level Q/C was performed by FastQC (v0.10.1). Pair-end reads of cDNA sequences were aligned back to the human genome (UCSC hg19 from Illumina iGenome) by the spliced read mapper TopHat (v2.0.4) with default option with “--mate-innder-dist 160” based on known transcripts of Ensembl Build GRCh37.75. For stranded RiboZero samples, TopHat used “--library-type fr-firststrand” option. The alignment statistics and Q/C were achieved by samtools (v0.1.18) and RSeQC (v2.3.5) to calculate quality control metrics on the resulting aligned reads, which provides useful information on mappability, uniformity of gene body coverage, insert length distributions and junction annotation, respectively. To achieve a gene-level expression profile, the properly paired and mapped reads were achieved by “samtools sort –n” option, and these reads were counted by htseq-count v0.5.3 (with intersection-strict mode and stranded option for RiboZero samples) according to gene annotation (Illumina iGenome), and RPKM was calculated. This provided 23,368 gene-level expression profiles.

### Statistics for SOX21 protein level

To determine the effect of a cell line of origin on nuclear SOX21 protein levels in Figure 1B, we used a mixed model comparing the mean expression levels across cell lines while accounting for the correlation of expression levels within replicate experiments (a total of 5 independent experiments were conducted) with a random intercept, implemented in R using the lme4 library and the lmer() function: expression~ as.factor (line)*condition*day+(1| replicate). To test the effect of the line of origin, this model was compared to a second model with no line effect using anova().

### Bioinformatic analyses

Principle component analysis was done using the prcomp() function in R. Agglomerative hierarchical clustering of genes using gene-level RPKM from RNA-seq data was performed using hclust() and cutree() with correlational distance (dist=1-r) in R. GWCoGAPS was run using default parameters as previously described (36, 37, 108, 109), for a range of k patterns (k=22 selected) and uncertainty as 10% of the data. Briefly, whole transcriptomic data was parallelized into seven sets. GWCoGAPS decomposes a matrix of experimental observations, **D**—here, log2 RNA-seq RPKMs—with genes as rows and samples as columns, into two matrices, by the following equation.

**D** ~ N(**AP**,Σ)

Where, **A** is the amplitude matrix indicating the strength of involvement of a given gene in each pattern, and **P** is the pattern matrix defining relationships (i.e., patterns) between samples. N and Σ are functions of each element of **AP** and represent the Normal distribution and the standard deviation, respectively. Projection of principal components and GWCoGAPS gene weights defines patterns of relationships between samples in new data associated with the gene expression signatures of the patterns from the primary data. These were achieved using the default projectR function in the projectR package as previously described (109). Enrichment was calculated via either the calcCoGAPSStat function in the CoGAPS Bioconductor package or the geneSetTest function in the limma Bioconductor package in R. ANOVAs were used to assess the association of each GWCoGAPS pattern with treatment, time, and cell line of origin, using lm() and summary() in R: lm(pattern~treatment*day+line).

### Gene age estimation

Gene age was estimated by phylostratigraphy that uses protein sequence similarity scored by BLASTP to find the minimal evolutionary age of protein-coding genes (57, 58). For each protein, the National Center for Biotechnology Information (NCBI) nonredundant database was used to find the most distant species in which a sufficiently similar protein sequence exists. We estimated the minimal evolutionary age of a gene as the age of the ancestor of the query species, human in this study, and the most distant species harboring a sufficiently similar sequence. To find the most distant species, we used the NCBI taxonomy for every species and estimated the timing of lineage divergence events with TimeTree (110). As in other studies, we used the e-value threshold of 10^−3^ to detect sequence similarity by BLASTP (58, 111, 112). For all human protein sequences, we filtered the sequences for a minimal length of 40 amino acids and a maximal length of 4,000 amino acids and kept only one protein isoform per gene (the longest and evolutionary oldest). We counted the number of genes in each phylostratum (PS), from the most ancient (PS 1, Cellular organisms) to the most recent (PS 31, *Homo sapiens*). We aggregated gene counts from individual phylostrata into five broad evolutionary eras: Ancient (PS 1 to 3, Cellular organisms to Opisthokonta; 4290 to 2101 millions of years ago (MYA)); Animal (PS 4 to 7, Metazoa to Deuterostomia, 2101 to 747 MYA); Chordate (PS 8 to 17, Chordate to Amniota, 747 to 320 MYA); Mammal (PS 18 to 22, Mammalia to Euarchontoglires, 320 to 91 MYA); Primate (PS 23 to 31, Primates to *Homo sapiens*, 91 Mya to present).

### ChIP-seq

The cells were crosslinked in 1% formaldehyde for 10 min at room temperature with constant agitation, followed by quenching with 125 mM glycine for 5 min. Nuclei were collected on days 2, 4, and 6 in self-renewal. Chromatin was fragmented with micrococcal nuclease (MNase) until the majority of DNA was in the range of 200-700 base pairs. Chromatin was incubated with an H3K9me3 antibody (13969, Cell signaling) at 4°C overnight, with constant agitation. Antibody-bound chromatin was immunoprecipitated with Protein G Dynabeads for 1 hour at 4°C, with constant agitation. Protein-DNA complexes were eluted from Dynabeads, and the DNA was reverse crosslinked with 0.3 M NaCl with 4 h incubation at 65°C. Free DNA was purified with QIAquick Gel Extraction Kit (28706X4, Qiagen). ChIP-seq libraries were generated using TruSeq ChIP Library Preparation Kit (Illumina, IP-202-1012) and the libraries were sequenced on Illumina HiSeq 3000. ChIP-seq raw reads were aligned to the human genome assembly hg19 using Burrows-Wheeler Aligner (BWA), and peak calling was done using MACS2.1.1. To understand H3K9me3 occupancy in promotors, ChIP-seq reads within +/−3kb of TSS were counted using bedtools.

## Author Contributions

S.K., S.S., G.SO., A.J., J.G.C., D.J.H., C.C., and R.D.M. conceived the study. S.K., S.S., and Y.W. performed cell culture and differentiation. Y.W. generated iPSC lines. K.O. generated SOX21-KO lines. S.K., A.J., Y.W., and K.O generated RNA-seq data. S.S. generated ChIP-seq data. G.SO., S.S., E.J.F., C.C., and J.S. developed and applied informatics methods to analyze RNA-seq data. V.L. and A.K. performed evolutionary gene age analysis. S.S. and C.C. analyzed ChIP-seq data. T.M.H., J.K., and D.R.W. provided fibroblasts for iPSC line generation and human brain tissue data. S.K., S.S., and N.M. performed immunocytochemistry. S.K., S.S., A.J., T.V., and D.J.H. performed high-content image analysis. S.K., S.S., G.SO., N.M, J.G.C., D.J.H., N.S., C.C., and R.D.M. interpreted the data. N.S., R.B., A.J.C., N.J.B., D.R.W., and R.D.M. directed the research. S.K., S.S., C.C., and R.D.M. wrote the manuscript. All authors participated in the discussion of results and manuscript editing.

## Acknowledgments

We thank the Lieber and Maltz families for their generous support of this work at the Lieber Institute for Brain Development (LIBD). This work was also supported by NCI/NIH grants R01CA177669, P30CA006973, U01CA212007, and U01CA253403 and the Johns Hopkins University Catalyst Award awarded to E.J.F., and R01NS116418, R01HG010898, MH116488, and U01MH124619 awarded to N.S. Data sharing and visualization via NeMO Analytics were supported by grants R24MH114815 and R01DC019370. We thank J. Jessee, MTI-GlobalStem, for the technical support of CRISPR/Cas9 plasmid transfection. We thank many members of LIBD and the Sestan lab for their helpful comments on this work.

## Supplementary Figures

**Fig. S1.**
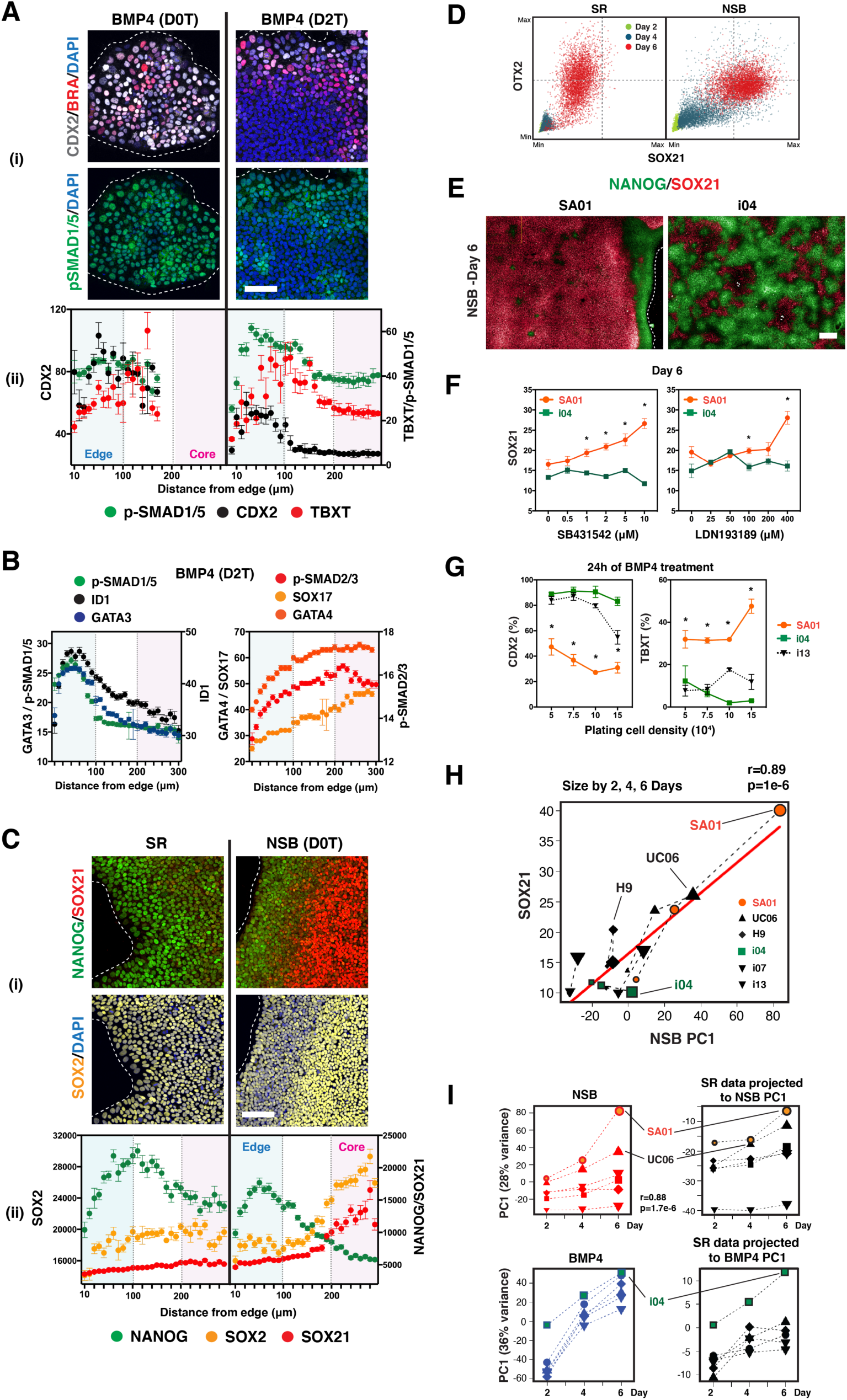
Cell line variation in the emergence of neural fate from pluripotency. (**A**) CDX2, TBXT, and pSMAD1/5 levels at 24 hours after BMP4 treatments (D0T: treated on day 0, D2T: treated on day 2). (i) Representative images. Scale bar, 100 μm. (ii) Expression levels plotted against distance from the edge. (**B**) Spatial expression of mesendoderm regulators at 24 h after BMP4 treatment on day 2 (D2T). (**C**) Spatial expression of NANOG, SOX21, and SOX2 on day 4 in SR and NSB. (i) Representative images. Scale bar, 100 μm. (ii) Spatial expression levels. (**D**) Scatter plots showing single-cell levels of SOX21 and OTX2 expression in self-renewal (SR) and NSB conditions through culture time demonstrate a positive correlation between them during neuroectodermal differentiation. (**E**) SOX21 and NANOG expression on day 6 of NSB treatment in SA01 and i04 lines show a more efficient formation of the core zone in SA01 compared to i04. Scale bar, 200 μm. Dashed lines indicate the edge of colonies. (**F**) Dose-response curve of BMP/TGFβ signaling inhibitors LDN193189 and SB431542 on SOX21 induction on day 6 shows differential responses between SA01 and i04 lines. *, p<0.05 between lines. (**G**) Proportion of CDX2 and BRACHURY (TBXT) expressing cells in SA01 and i04 lines after 24 h of BMP4 treatment on day 0 (D0T) show the cell line difference is not caused by initial cell plating density. *, p<0.05 between lines. (**H**) Scatter plot showing the correlation between SOX21 protein levels in NSB (Fig. 1B) and NSB PC1. (**I**) Projections of SR data into PC1 of data from differentiation showing line-specific transcriptional lineage priming. PC1 of NSB and BMP4 data (left) and projection of SR data into the NSB and BMP4 PC1 (right).

**Fig. S2.**
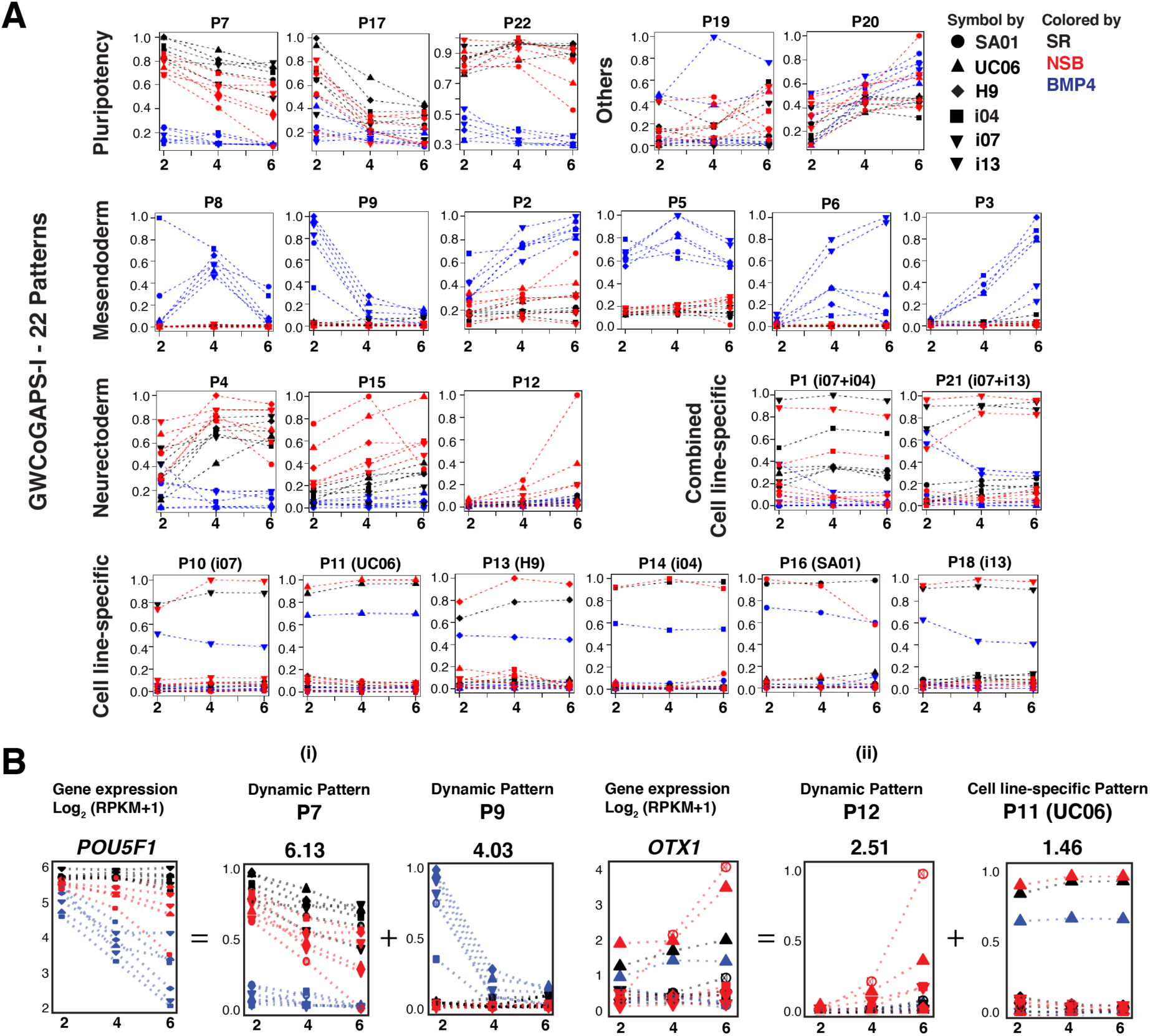
Dynamic and stable transcription modules defined by GWCoGAPS-I. (**A**) The 22 patterns generated by GWCoGAPS decomposition across 6 cell lines, 3 conditions, and 3 times. Patterns are presented in groups of similar characteristics; dynamic patterns (pluripotency, BMP4-response mesendoderm, NSB-response neuroectoderm, and others) or cell line-specific patterns (6 cell line-specific patterns and 2 combined cell line-specific patterns). (**B**) The use of each GWCoGAPS pattern across genes can be precisely defined by gene-specific amplitudes for all patterns (Supplementary Table S2). Two examples of individual genes whose expression patterns are represented by the combination of multiple GWCoGAPS patterns are shown. (i) The complete expression of *POU5F1* is represented by two dynamic GWCoGAPS patterns P7 and P9. (ii) The complete expression of *OTX1* is represented by a dynamic GWCoGAPS pattern P12 and the UC06 line-specific GWCoGAPS pattern P11.

**Fig. S3.**
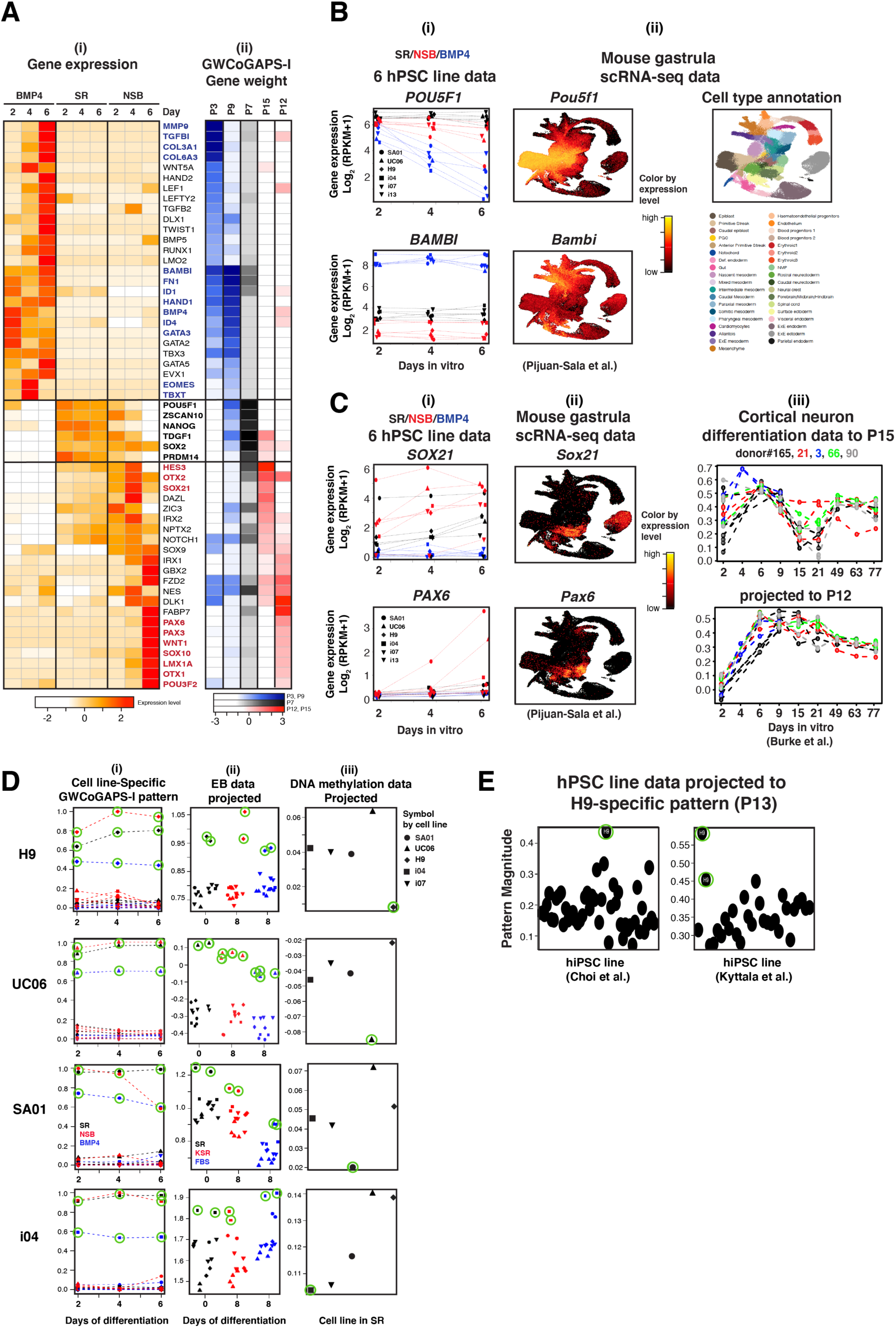
Decomposing dynamic and stable transcription modules. (**A**) Gene expression of pluripotency or differentiation regulators and their weights in dynamic GWCoGAPS patterns. (i) Heatmap of gene expression. (ii) Heatmap showing the gene weights. SR pattern: P7; NSB pattern: P12 and P15; BMP4 pattern: P9 and P3. (**B**) GWCoGAPS pattern P7 represents the loss of pluripotency and P3 represents the induction of mesendoderm regulators. (i) Expression of *POU5f1* and *BAMBI*, the top-ranked genes in P7 and P3, respectively, in 6 hPSC line data. (ii) Expression of *Pou5f1* and *Bambi* in mouse gastrula scRNA-seq data. (**C**) NSB patterns P15 and P12 represent distinct transition steps toward neural fates. (i) Expression of *SOX21* and *PAX6*, the top-ranked genes in P15 and P12, respectively, in 6 hPSC line data. (ii) Expression of *Sox21* and *Pax6* in mouse gastrula scRNA-seq data. (iii) Projection of cortical neuron differentiation data from multiple hiPSC lines. (**D**) Projections of microarray and DNA methylation data generated from multiple hPSC lines into the cell line-specific GWCoGAPS-I patterns demonstrate the stability of the cell line-specific transcriptional signatures within a cell line. (i) H9, UC06, SA01, and i04 line-specific GWCoGAPS-I patterns that define distinct transcriptional signatures from all other lines across time and condition. Each cell line sample is circled in green. (ii) Projections of microarray dataset containing the same 6 lines under embryoid body (EB) differentiation conditions (SR for self-renewal in ES medium plus FGF2, KSR for ectodermal differentiation, and FBS for mesendodermal differentiation) into each cell line-specific patterns discriminate the corresponding cell line samples from all other lines. (iii) Projections of DNA methylation data show that promoters of genes expressed specifically in each cell line are hypomethylated in the corresponding cell line. (**E**) Projection of multiple hPSC line data into H9-specific pattern. H9 samples are circled in green.

**Fig. S4.**
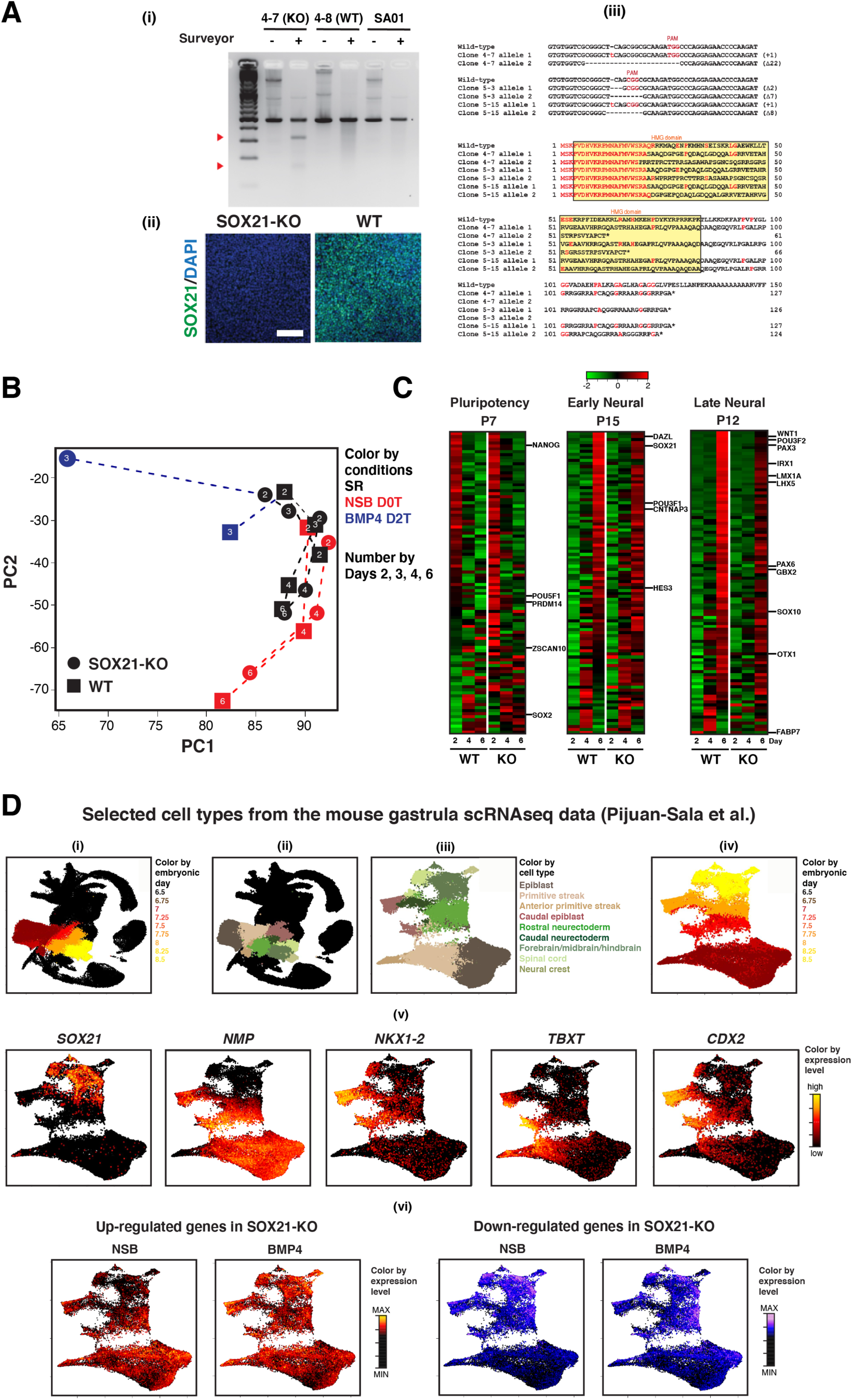
SOX21 regulates early forebrain fate choice. (**A**) Establishment of SOX21-KO lines by CRISPR/Cas9 technology. All SOX21-KO SA01 hPSC lines were screened using Surveyor and immunofluorescence assays and verified by DNA sequencing. (i) Analysis of SOX21-KO clones using Surveyor assay. The gel image shows modification at the SOX21 locus in a clone 4-7. Red arrowheads indicate expected fragment sizes for the SOX21 locus. (ii) Immunostaining of WT and SOX21-KO clones shows complete loss of SOX21 expression in clone 4-7. The cells were cultured in the presence of NSB for 6 days. Scale bar, 100 μm. (iii) Amino acid sequence of SOX21 alterations by CRISPR/Cas9 confirms knockout of SOX21. In three clones (clones 4-7, 5-3, and 5-15), frameshift mutation, premature stop codon mutation, or mutation that disrupts the HMG domain were confirmed in both *SOX21* alleles. These three clones were used for the functional assays shown in Fig. 3. (**B**) Projection of WT and SOX21-KO line data into PC1 and PC2 of days 2, 4, and 6 data (Fig. 1C) reveals delayed neuroectodermal differentiation in NSB and accelerated early mesendodermal differentiation under BMP4 D2T in SOX21-KO compared to WT. (**C**) Heatmaps showing the top 100 genes from P7, P15, and P12 in NSB. (**D**) Uniform manifold approximation and projection (UMAP) plots of selected epiblast and neural cell populations in mouse gastrula RNA-seq data showing *SOX21* and neuromesodermal precursor (NMP) gene expression. (i) A UMAP plot showing all cells with selected cell types colored by embryonic days. (ii) A UMAP plot showing all cells colored by cell type annotation with selected cell types (iii) A UMAP plot showing the selected cell types only and cell type annotation. (iv) A UMAP plot showing the selected cell types colored by embryonic days. (v) Expression of *Sox21* and average expression of the neuromesodermal progenitor (NMP) genes in the epiblast and early neural lineage cells in mouse gastrula RNA-seq data. *Nkx1-2* expression delineates caudal epiblast cells showing the NMP gene signature, while *Tbxt* expression represents both the caudal epiblast cells and anterior primitive streak cells. Note the mutually exclusive expression between *Sox21* in rostral neuroectoderm and *Cdx2* in caudal epiblast at developmental day 7.75, showing the early role of SOX21 in anterior-posterior axis patterning. (vi) A UMAP plot showing the projection of log fold changes of up- or down-regulated genes from the SOX21-KO RNA-seq data into the mouse gastrula RNA-seq data. Color scales range from the maximum to the minimum of the projected values within an analysis for each panel. The projected values are comparable within a panel but not between panels.

**Fig. S5.**
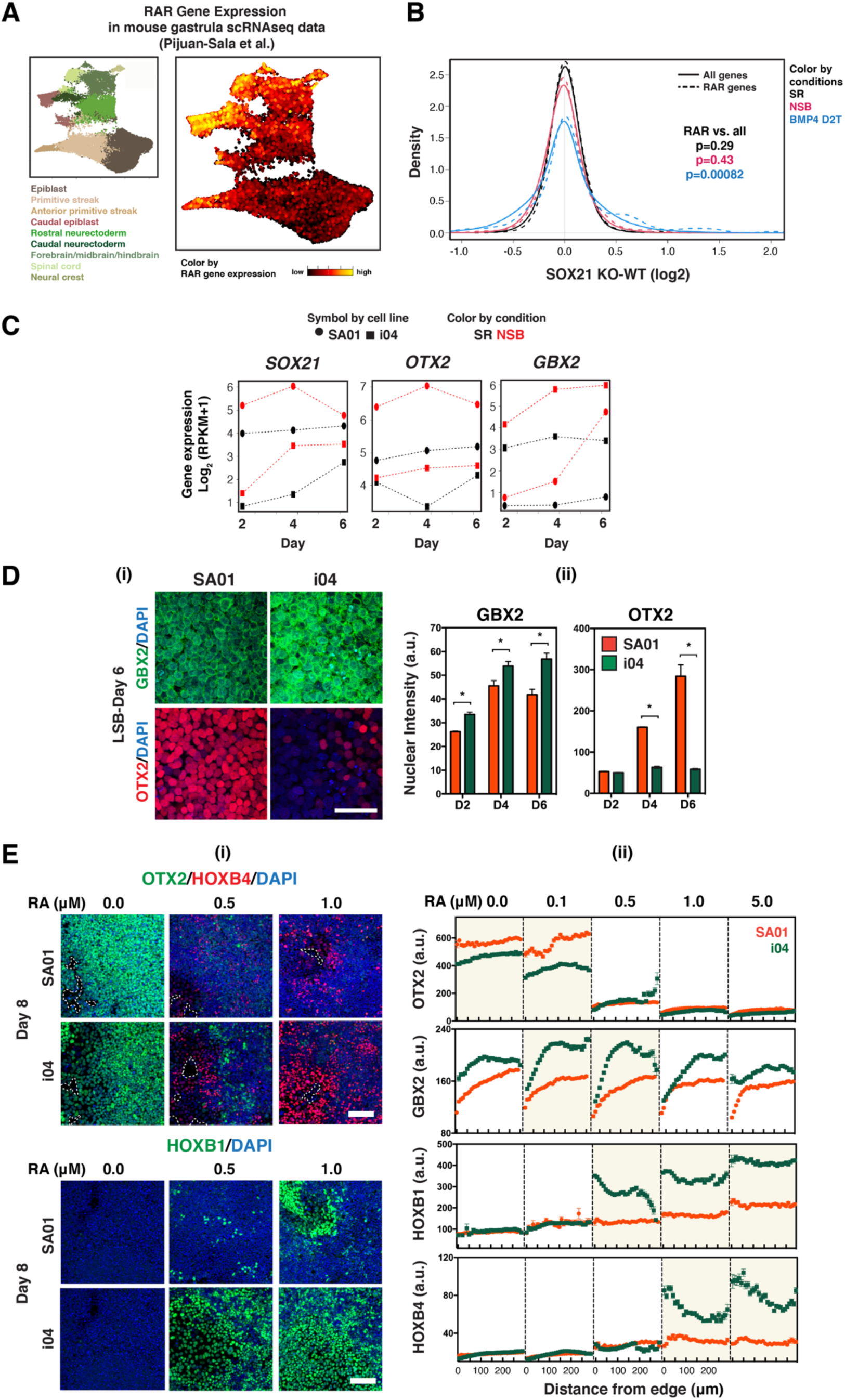
Cell line variation in fore- vs. hind-brain fate bias. (**A**) A UMAP of the selected populations from the mouse gastrula scRNA-seq data colored by retinoic acid responsive (RAR) gene expression. (**B**) A density plot of all 252 RAR genes (dotted lines) compared to all 21,022 genes (solid lines) in SR, NSB, and BMP4 D2T conditions of SOX21-KO RNA-seq data. (**C**) mRNA expression of *OTX2*, *GBX2*, and *SOX21* from the hiPSC RNA-seq data. (**D**) Differential GBX2 and OTX2 expression in SA01 and i04. (i) Representative images on day 6 in neuroectoderm differentiation conditions (LDN193189 and SB431542, LSB). Scale bar, 50 μm. (ii) OTX2 and GBX2 expression level in SR. *, Comparison between SA01 and i04 (p<0.05). (**E**) Dose-response analysis of RA shows differential posterization of neural precursors between SA01 and i04 lines. (i) Representative images show differential OTX2, HOXB4, and HOXB1 expression on day 8 of LSB+RA-induced differentiation between SA01 and i04 lines. Scale bar, 100 μm. Dashed lines indicate the edge of colonies. (ii) Dose-response analysis of RA on spatial expression of anterior-posterior neural regulators shows a sequential posteriorization from the core to edge zones and a cell line-specific RA response. Single-cell levels of OTX2, GBX2, HOXB1, and HOXB4 expression are measured on day 8 of LSB+RA-induced differentiation and plotted against the distance from the edge. In contrast, HOXB9 and HOXD9, markers of the more posterior spinal cord and trunk neural crest, were not detected in RA-treated cells (data not shown).

**Fig. S6.**
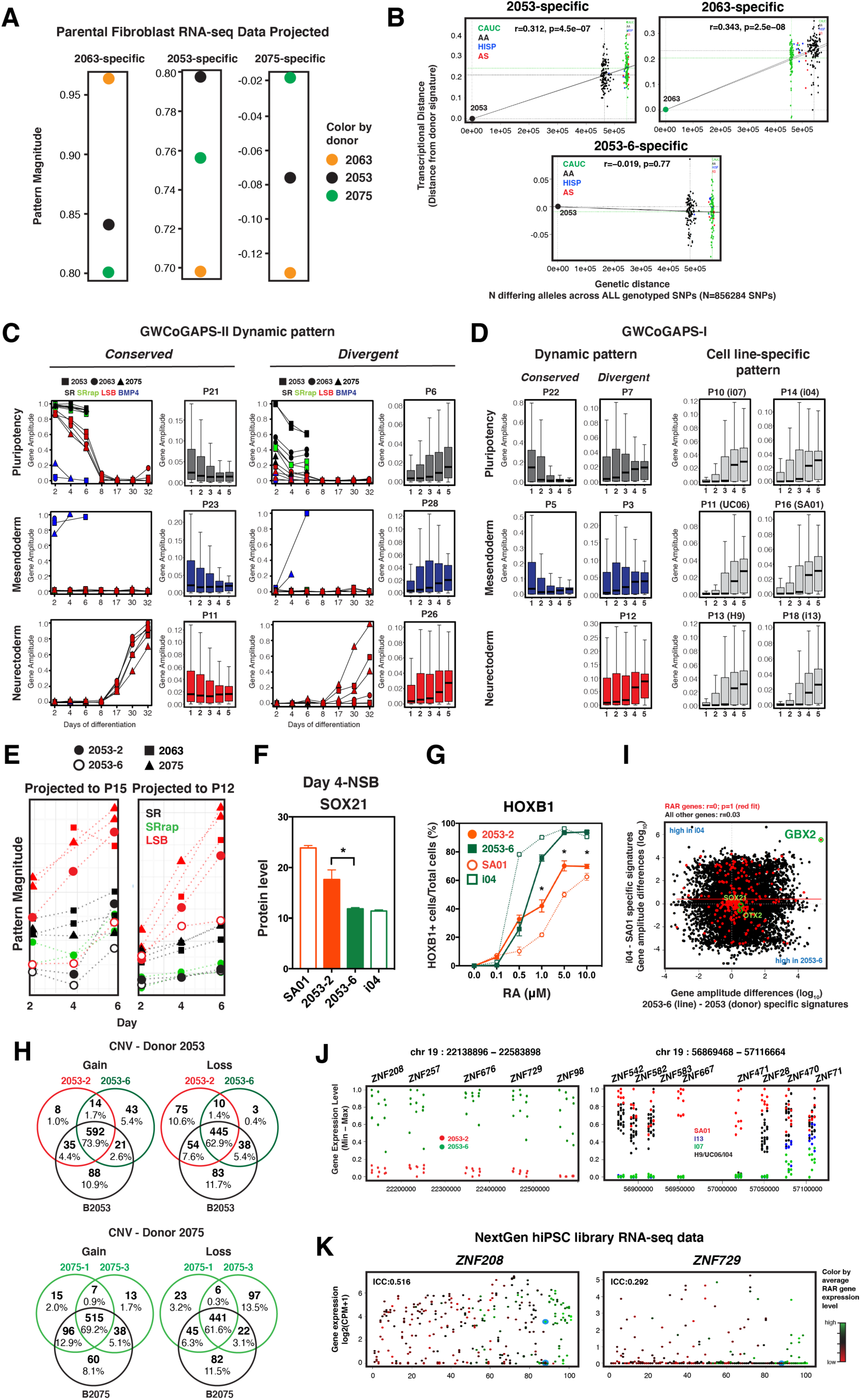
Genetic and epigenetic elements contribute to the donor- and line-specific transcriptional signatures. (**A**) Projection of parental fibroblast RNA-seq data into donor-specific patterns shows donor-specific transcriptional signatures are highest in the parental fibroblasts of the corresponding donor. (**B**) Scatter plots showing the association between genetic similarity and the strength of donor-specific transcription signatures across pair-wise comparisons with other donors. Each point is a comparison of the highlighted donor to another donor. Genetic distance between donors was quantified as the number of differing alleles across all genotyped single nucleotide polymorphisms (SNPs on the X-axis). The transcriptional distance was quantified as the strength of the donor-specific transcriptional signature in other donors’ tissue (Y-axis; same data depicted in Y-axis of Fig. 6Aii and 6Cii, centered such that the source donor has a value of zero). Pearson’s correlation and p-values were calculated by omitting the donor data point at the origin. Solid black lines pass through the origin and each ethnicity’s average Y-value, indicated as horizontal dashed lines. Ethnicity of donors is indicated by color: CAUC, Caucasian; AA, African American; HISP, Hispanic; AS, Asian. (**C**) Contribution of genes of 5 evolutionary eras (1, Ancient; 2, Animal; 3, Chordate; 4, Mammal; 5, Primate) to the conserved or divergent dynamic patterns of GWCoGAPS-II analysis. Dynamic patterns conserved across cell lines show high gene amplitudes in ancient genes, while dynamic patterns divergent across cell lines show higher gene amplitudes in newer genes. Comparison on the distribution of era 1 gene amplitudes to era 5 gene amplitudes by Wilcoxon rank sum test: p>1e-16 for all patterns shown. (**D**) Contribution of genes of 5 evolutionary eras to the conserved or divergent dynamic and cell line-specific patterns of GWCoGAPS-I analysis (p<1e-16 for all patterns shown). (**E**) Projection of RNA-seq data into the neural patterns P15 and P12. (**F**) Differential SOX21 levels in SA01, i04, and two 2053 lines. *, Comparison between 2053-2 and 2053-6 (p<0.05). (**G**) Two replicate lines from donor 2053 show differential responsiveness to RA. Number of HOXB1 expressing cells on day 8 in response to varying doses of RA in SA01, i04, and two 2053 lines. *, Comparison between 2053-2 and 2053-6 (p<0.05). (**H**) The Venn diagrams showing the copy number variants (CNVs) gained and lost in the two replicate iPSC lines and the mature post-mortem brain tissue of donors 2053 and 2075 with respect to the reference genome. In all cases, the vast majority of differences from the reference genome are the same across both iPSC lines and the brain tissues derived from each donor, indicating little change in genomic DNA. A similar analysis for single nucleotide variants (SNVs) showed similar results (data not shown). (**I**) Scatter plot comparing differences in gene amplitudes between the i04 and SA01 line-specific patterns with differences in gene amplitudes between 2053 donor-specific and 2053-6 line-specific patterns. (**J**) Differential expression of the clustered ZNF genes across the discordant iPSC lines 2053-2 and 2053-6 (left) and across the 6 hPSC lines (right). Plots show expression of each gene from its minimum to its maximum in pluripotency, NSB/LSB, and BMP4 conditions for all lines. (**K**) Expression of ZNF208 and ZNF729, two of the genes expressed specifically in line 2053-6 (Supplementary Fig. S6*J*) in the NextGen Consortium RNA-seq data. Replicate lines from individual donors are displayed as points along a single X-axis value. Donors are ordered by their rank in the first principal component of this data and colored by RAR gene expression (see Fig. 6*A* and Supplementary Table S5 and S7 for more detail on this data set). ZNF208 shows high expression in most lines, with a minority of lines showing very low expression. In contrast, ZNF729 showed no reads in most lines, and a few lines showed significant expression. While quite distinct from one another, the expression patterns of both genes are consistent with epigenetic regulation, i.e., divergent expression levels across multiple lines derived from the same donor. Expression of these and other genes can be explored at: https://nemoanalytics.org/p?l=Kim2023&g=ZNF208. ICC= intra-class correlation coefficient (i.e., the proportion of variance that can be accounted for by donor).

**Fig. S7.**
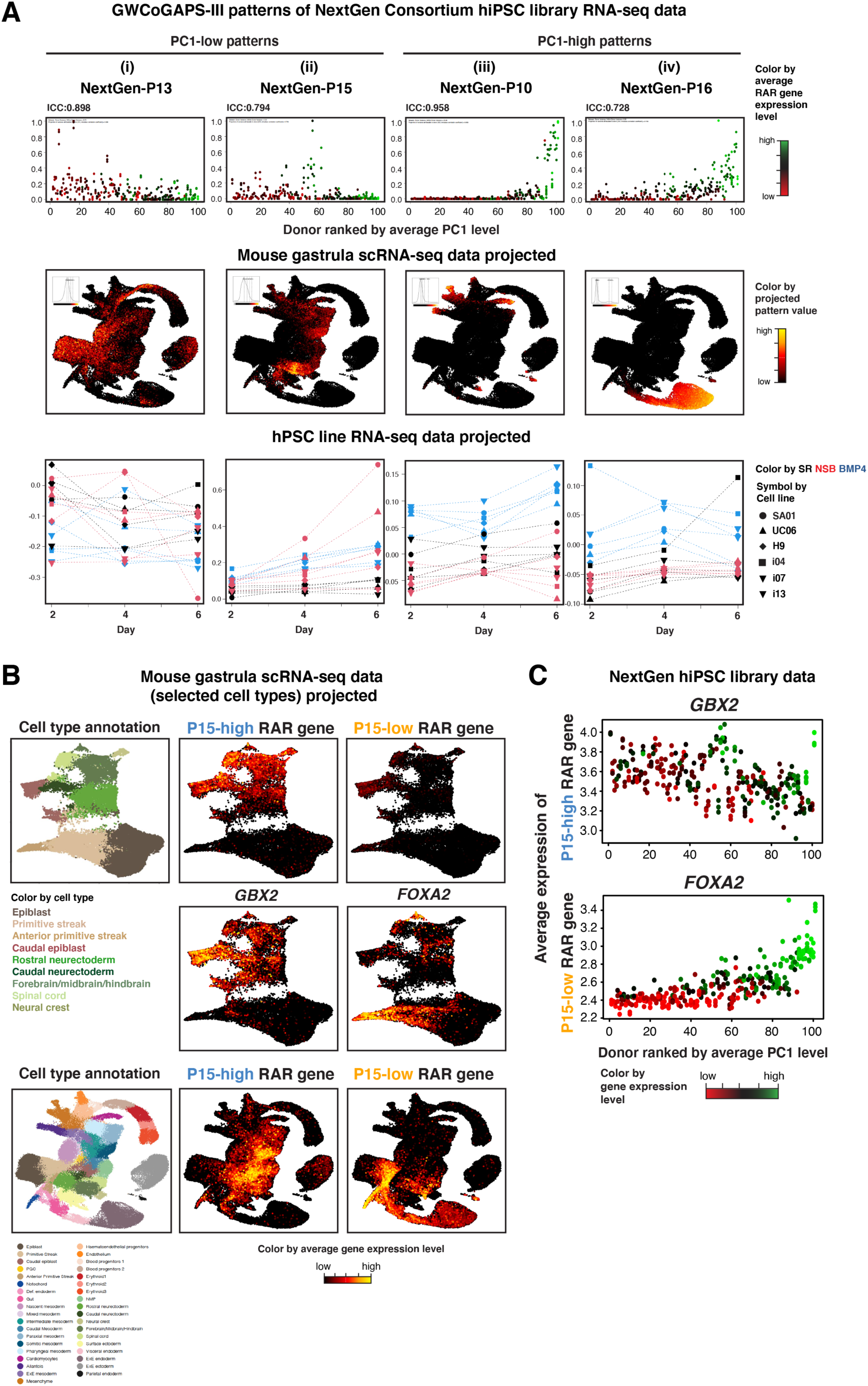
Early developmental bias and RA signaling define hPSC variation in the wider human population. (**A**) 4 of 20 GWCoGAPS-III patterns from the NextGen RNA-seq data and their projection into the mouse gastrula RNA-seq data and the hPSC RNA-seq data (this study), showing that the low and high PC1 signals each correspond to multiple distinct lineage elements: low PC1 corresponds to pluripotency and the emerging neural lineages (i, ii), while high PC1 corresponds to mesodermal and endodermal fates (iii, iv). (**B**) A UMAP of the mouse gastrula scRNA-seq data colored by expression of RAR genes with high or low P15 gene amplitudes and *GBX2* and *FOXA2* expression. (**C**) Average expression level of RAR genes with high or low P15 gene amplitudes in NextGen RNA-seq data colored by *GBX2* and *FOXA2* expression.

## Supplementary Table Legends

**Supplementary Table S1. Mapping and annotation details of RNA-seq and summary of experimental times and conditions for samples**

**Supplementary Table S2. Individual gene amplitudes for each of the 22 GWCoGAPS-I patterns. All 22 patterns are presented in Supplementary Fig. S2*A***.

**Supplementary Table S3. List of top 100 genes in the 22 GWCoGAPS-I patterns.**

**Supplementary Table S4. Individual gene amplitudes for GWCoGAPS-II. Donor- and line-specific patterns (P4, P17, P20, and P27) are defined by GWCoGAPS-II.**

**Supplementary Table S5. Genes ranked by their contribution to the line- and donor-specific patterns.** Sheet1: GWCoGAPS-I genes, Sheet2: GWCoGAPS-II genes. Submitting the top 250 genes from these lists to the Enrichr webtool (https://maayanlab.cloud/Enrichr/enrich?dataset=3f14ebb0f1889f3a5acb6d940326bebb; https://maayanlab.cloud/Enrichr/enrich?dataset=cb12b784682d4aed17f5b17d6cdbee3b) revealed enrichment in 1) Co-expression with many KRAB-ZNF genes (p=7.8e-9 to p=5.8e-6) from the ARCHS4 database, 2) H3K9me3 (p=1.8e-19 to p=6.4e-7) and SETDB1 (p=9.7e-8) ChIP-seq peaks from the ENCODE Histone modifications and TF ChIP-seq databases, and 3) TRIM28/KAP1 protein-protein interactions from the BioPlex database (p=6.8e-6). These enrichments represent a canonical mechanism of transcriptional repression: KRAB-ZNF genes bind repetitive genomic DNA derived from TEs and TRIM28/KAP1, recruiting SETDB1 to deposit the H3K9me3. We also found enrichment of protocadherin (PCDH) genes among the GWCoGAPS gene amplitudes in line-and donor-specific patterns (p=2e-17 and p=0.00046, in GWCoGAPS-I and –II, respectively). Y chromosome genes in these lists serve as a control positive in that their expression patterns are described primarily as a combination of the male line-specific patterns. Genes in line- and donor-specific expression patterns can be explored in our NeMO Analytics portal, where examples of both epigenetic (e.g., ZNF genes in Supplementary Fig. S6J and S6K) and genetic (e.g., NOMO3) regulation can be observed in our hPSC lines and the NextGen Consortium lines: https://nemoanalytics.org/p?l=Kim2023&g=NOMO3.

**Supplementary Table S6. Normalized H3K9me3 levels in 2053-2 and 2053-6 lines in self renewal**

**Supplementary Table S7. Values of retinoic acid response (RAR) genes in PC1 of NextGen hiPSC library RNAseq data and GWCoGAPS-I patterns 14, 15, and 16.** P14, i04 line-specific pattern; P15, neuroectoderm pattern; P16, SA01 line-specific pattern.

**Supplementary Table S8. Individual gene amplitudes for NextGen GWCoGAPS-III.**

**Supplementary Table S9. Differentially expressed genes between two groups of hiPSC lines with neural bias.**

## Notes

https://nemoanalytics.org/p?l=Kim2023&g=GBX2

### Competing Interest Statement

The authors have declared no competing interest.

### Summary of Updates

The manuscript revised with new findings and additional data.

## References

1. S. J. Arnold, E. J. Robertson, Making a commitment: cell lineage allocation and axis patterning in the early mouse embryo. Nature reviews. Molecular cell biology 10, 91–103 (2009).

2. E. M. De Robertis, Spemann’s organizer and the self-regulation of embryonic fields. Mechanisms of development 126, 925–941 (2009).

3. I. G. Brons et al., Derivation of pluripotent epiblast stem cells from mammalian embryos. Nature 448, 191–195 (2007).

4. P. J. Tesar et al., New cell lines from mouse epiblast share defining features with human embryonic stem cells. Nature 448, 196–199 (2007).

5. G. Guo et al., Naive Pluripotent Stem Cells Derived Directly from Isolated Cells of the Human Inner Cell Mass. Stem Cell Reports 6, 437–446 (2016).

6. S. Bao et al., Epigenetic reversion of post-implantation epiblast to pluripotent embryonic stem cells. Nature 461, 1292–1295 (2009).

7. S. Temple, L. Studer, Lessons Learned from Pioneering Neural Stem Cell Studies. Stem Cell Reports 8, 191–193 (2017).

8. J. Choi et al., A comparison of genetically matched cell lines reveals the equivalence of human iPSCs and ESCs. Nature biotechnology 33, 1173–1181 (2015).

9. F. Rouhani et al., Genetic background drives transcriptional variation in human induced pluripotent stem cells. PLoS genetics 10, e1004432 (2014).

10. C. DeBoever et al., Large-Scale Profiling Reveals the Influence of Genetic Variation on Gene Expression in Human Induced Pluripotent Stem Cells. Cell stem cell 20, 533–546 e537 (2017).

11. H. Kilpinen et al., Common genetic variation drives molecular heterogeneity in human iPSCs. Nature 546, 370–375 (2017).

12. A. S. E. Cuomo et al., Single-cell RNA-sequencing of differentiating iPS cells reveals dynamic genetic effects on gene expression. Nature Communications 11, 810 (2020).

13. I. Carcamo-Orive et al., Analysis of Transcriptional Variability in a Large Human iPSC Library Reveals Genetic and Non-genetic Determinants of Heterogeneity. Cell stem cell 20, 518–532.e519 (2017).

14. A. Strano, E. Tuck, V. E. Stubbs, F. J. Livesey, Variable Outcomes in Neural Differentiation of Human PSCs Arise from Intrinsic Differences in Developmental Signaling Pathways. Cell reports 31, 107732 (2020).

15. N. Micali et al., Variation of Human Neural Stem Cells Generating Organizer States In Vitro before Committing to Cortical Excitatory or Inhibitory Neuronal Fates. Cell reports 31, 107599 (2020).

16. J. Mariani et al., FOXG1-Dependent Dysregulation of GABA/Glutamate Neuron Differentiation in Autism Spectrum Disorders. Cell 162, 375–390 (2015).

17. E. E. Burke et al., Dissecting transcriptomic signatures of neuronal differentiation and maturation using iPSCs. Nature communications 11, 462 (2020).

18. S. Kanton et al., Organoid single-cell genomic atlas uncovers human-specific features of brain development. Nature 574, 418–422 (2019).

19. M. Wang et al., Increased Neural Progenitor Proliferation in a hiPSC Model of Autism Induces Replication Stress-Associated Genome Instability. Cell stem cell 26, 221–233.e226. (2020).

20. A. Jourdon et al., ASD modelling in organoids reveals imbalance of excitatory cortical neuron subtypes during early neurogenesis. bioRxiv 10.1101/2022.03.19.484988, 2022.2003.2019.484988 (2022).

21. B. Paulsen et al., Autism genes converge on asynchronous development of shared neuron classes. Nature 10.1038/s41586-021-04358-6 (2022).

22. J. Orvis et al., gEAR: Gene Expression Analysis Resource portal for community-driven, multi-omic data exploration. Nature methods 18, 843–844 (2021).

23. Shelley R. Hough et al., Single-Cell Gene Expression Profiles Define Self-Renewing, Pluripotent, and Lineage Primed States of Human Pluripotent Stem Cells. Stem Cell Reports 22, 881–895 (2014).

24. G. Guo et al., Human naive epiblast cells possess unrestricted lineage potential. Cell stem cell 10.1016/j.stem.2021.02.025 (2021).

25. M. Nakanishi et al., Human Pluripotency Is Initiated and Preserved by a Unique Subset of Founder Cells. Cell 177, 1–15 (2019).

26. A. Warmflash, B. Sorre, F. Etoc, E. D. Siggia, A. H. Brivanlou, A method to recapitulate early embryonic spatial patterning in human embryonic stem cells. Nature methods 10.1038/nmeth.3016 (2014).

27. F. Etoc et al., A Balance between Secreted Inhibitors and Edge Sensing Controls Gastruloid Self-Organization. Developmental cell 39, 302–315 (2016).

28. P. Rifes et al., Modeling neural tube development by differentiation of human embryonic stem cells in a microfluidic WNT gradient. Nature biotechnology 38, 1265–1273 (2020).

29. S. M. Chambers et al., Highly efficient neural conversion of human ES and iPS cells by dual inhibition of SMAD signaling. Nature biotechnology 27, 275–280 (2009).

30. T. Faial et al., Brachyury and SMAD signalling collaboratively orchestrate distinct mesoderm and endoderm gene regulatory networks in differentiating human embryonic stem cells. Development 142, 2121–2135 (2015).

31. S. Mendjan et al., NANOG and CDX2 Pattern Distinct Subtypes of Human Mesoderm during Exit from Pluripotency. Cell stem cell 15, 310–325 (2014).

32. A. N. Kuzmichev et al., Sox2 acts through Sox21 to regulate transcription in pluripotent and differentiated cells. Current biology : CB 22, 1705–1710 (2012).

33. A. Simeone, E. Puelles, D. Acampora, The Otx family. Curr Opin Genet Dev 12, 409–415 (2002).

34. B. S. Mallon et al., StemCellDB: the human pluripotent stem cell database at the National Institutes of Health. Stem cell research 10, 57–66 (2013).

35. G. Sharma, C. Colantuoni, L. A. Goff, E. J. Fertig, G. Stein-O’Brien, projectR: An R/Bioconductor package for transfer learning via PCA, NMF, correlation, and clustering. Bioinformatics 36, 3592–3593 (2020).

36. E. J. D. Fertig, J., A. V. P. Favorov, G. Ochs, M. F., CoGAPS: an R/C++ package to identify patterns and biological process activity in transcriptomic data. Bioinformatics 26, 2792–2793 (2010).

37. G. L. Stein-O’Brien, et al., PatternMarkers & GWCoGAPS for novel data-driven biomarkers via whole transcriptome NMF. Bioinformatics 10.1093/bioinformatics/btx058, 1–3 (2017).

38. B. Pijuan-Sala et al., A single-cell molecular map of mouse gastrulation and early organogenesis. Nature 566, 490–495 (2019).

39. A. Kyttala et al., Genetic Variability Overrides the Impact of Parental Cell Type and Determines iPSC Differentiation Potential. Stem Cell Reports 6, 200–212 (2016).

40. I. Costello et al., The T-box transcription factor Eomesodermin acts upstream of Mesp1 to specify cardiac mesoderm during mouse gastrulation. Nature cell biology 13, 1084–1091 (2011).

41. D. Henrique, E. Abranches, L. Verrier, K. G. Storey, Neuromesodermal progenitors and the making of the spinal cord. Development 142, 2864–2875 (2015).

42. C. Guibentif et al., Diverse Routes toward Early Somites in the Mouse Embryo. Developmental cell 56, 141–153.e146 (2020).

43. A. Gogolou et al., Early anteroposterior regionalisation of human neural crest is shaped by a pro-mesodermal factor. eLife 11 (2022).

44. N. Hu, P. H. Strobl-Mazzulla, M. Simoes-Costa, E. Sanchez-Vasquez, M. E. Bronner, DNA methyltransferase 3B regulates duration of neural crest production via repression of Sox10. Proceedings of the National Academy of Sciences of the United States of America 111, 17911–17916 (2014).

45. N. Hu, P. Strobl-Mazzulla, T. Sauka-Spengler, M. E. Bronner, DNA methyltransferase3A as a molecular switch mediating the neural tube-to-neural crest fate transition. Genes & development 26, 2380–2385 (2012).

46. M. Goolam et al., Heterogeneity in Oct4 and Sox2 Targets Biases Cell Fate in 4-Cell Mouse Embryos. Cell 165, 61–74 (2016).

47. Z. Fang et al., SOX21 Ensures Rostral Forebrain Identity by Suppression of WNT8B during Neural Regionalization of Human Embryonic Stem Cells. Stem Cell Reports 13, 1–15 (2019).

48. S. Matsuda et al., Sox21 promotes hippocampal adult neurogenesis via the transcriptional repression of the Hes5 gene. The Journal of neuroscience : the official journal of the Society for Neuroscience 32, 12543–12557 (2012).

49. J. E. Balmer, Gene expression regulation by retinoic acid. The Journal of Lipid Research 43, 1773–1808 (2002).

50. N. B. Ghyselinck, G. Duester, Retinoic acid signaling pathways. Development 146 (2019).

51. M. Shibata et al., Regulation of prefrontal patterning and connectivity by retinoic acid. Nature 598, 483–488 (2021).

52. M. Gouti et al., A Gene Regulatory Network Balances Neural and Mesoderm Specification during Vertebrate Trunk Development. Developmental cell 41, 243–261 e247 (2017).

53. S. Millet et al., A role for Gbx2 in repression of Otx2 and positioning the mid/hindbrain organizer. Nature 401, 161–164 (1999).

54. R. Krumlauf, Hox Genes and the Hindbrain: A Study in Segments. Current topics in developmental biology 116, 581–596 (2016).

55. A. E. Jaffe et al., Developmental and genetic regulation of the human cortex transcriptome illuminate schizophrenia pathogenesis. Nature neuroscience 21, 1117–1125 (2018).

56. GTEx_Consortium, The Genotype-Tissue Expression (GTEx) pilot analysis: Multitissue gene regulation in humans. Science 348, 648–660 (2015).

57. T. Domazet-Loso, J. Brajkovic, D. Tautz, A phylostratigraphy approach to uncover the genomic history of major adaptations in metazoan lineages. Trends in genetics : TIG 23, 533–539 (2007).

58. J. A. Weber et al., The whale shark genome reveals how genomic and physiological properties scale with body size. Proceedings of the National Academy of Sciences of the United States of America 117, 20662–20671 (2020).

59. J. E. Ferrell, Jr., Bistability, bifurcations, and Waddington’s epigenetic landscape. Current biology : CB 22, R458–466 (2012).

60. R. L. Collins et al., A cross-disorder dosage sensitivity map of the human genome. Cell 185, 3041–3055 e3025 (2022).

61. E. B. Chuong, N. C. Elde, C. Feschotte, Regulatory activities of transposable elements: from conflicts to benefits. Nature reviews. Genetics 18, 71–86 (2017).

62. G. Ecco et al., Transposable Elements and Their KRAB-ZFP Controllers Regulate Gene Expression in Adult Tissues. Developmental cell 36, 611–623 (2016).

63. S. Lukic, J. C. Nicolas, A. J. Levine, The diversity of zinc-finger genes on human chromosome 19 provides an evolutionary mechanism for defense against inherited endogenous retroviruses. Cell death and differentiation 21, 381–387 (2014).

64. M. Florio et al., Evolution and cell-type specificity of human-specific genes preferentially expressed in progenitors of fetal neocortex. eLife 7 (2018).

65. J. Pontis et al., Hominoid-Specific Transposable Elements and KZFPs Facilitate Human Embryonic Genome Activation and Control Transcription in Naive Human ESCs. Cell stem cell 24, 724–735 e725 (2019).

66. D. Ortmann et al., Naive Pluripotent Stem Cells Exhibit Phenotypic Variability that Is Driven by Genetic Variation. Cell stem cell 27, 470–481.e476 (2020).

67. A. Vigilante et al., Identifying Extrinsic versus Intrinsic Drivers of Variation in Cell Behavior in Human iPSC Lines from Healthy Donors. Cell reports 26, 2078–2087 e2073 (2019).

68. M. J. Bonder et al., Identification of rare and common regulatory variants in pluripotent cells using population-scale transcriptomics. Nature genetics 53, 313–321 (2021).

69. J. Jerber et al., Population-scale single-cell RNA-seq profiling across dopaminergic neuron differentiation. Nature genetics 53, 304–312 (2021).

70. P. Puigdevall, J. Jerber, P. Danecek, S. Castellano, H. Kilpinen, Effects of somatic mutations on cellular differentiation in iPSC models of neurodevelopment. bioRxiv 10.1101/2022.03.04.482992, 2022.2003.2004.482992 (2022).

71. F. T. Merkle et al., Whole-genome analysis of human embryonic stem cells enables rational line selection based on genetic variation. Cell stem cell 29, 472–486 e477 (2022).

72. C. Bock et al., Reference Maps of human ES and iPS cell variation enable high-throughput characterization of pluripotent cell lines. Cell 144, 439–452 (2011).

73. B. Pijuan-Sala et al., Single-cell chromatin accessibility maps reveal regulatory programs driving early mouse organogenesis. Nature cell biology 22, 487–497 (2020).

74. S. Grosswendt et al., Epigenetic regulator function through mouse gastrulation. Nature 584, 102–108 (2020).

75. H. M. Chen et al., Transcripts involved in calcium signaling and telencephalic neuronal fate are altered in induced pluripotent stem cells from bipolar disorder patients. Translational psychiatry 4, e375 (2014).

76. J. M. Madison et al., Characterization of bipolar disorder patient-specific induced pluripotent stem cells from a family reveals neurodevelopmental and mRNA expression abnormalities. Molecular psychiatry 20, 703–717 (2015).

77. M. C. Marchetto et al., Altered proliferation and networks in neural cells derived from idiopathic autistic individuals. Molecular psychiatry 22, 820–835 (2017).

78. D. Rosebrock et al., Enhanced cortical neural stem cell identity through short SMAD and WNT inhibition in human cerebral organoids facilitates emergence of outer radial glial cells. Nature cell biology 24, 981–995 (2022).

79. W. R. Reay, M. J. Cairns, The role of the retinoids in schizophrenia: genomic and clinical perspectives. Molecular psychiatry 25, 706–718 (2020).

80. H. Yu et al., Dynamic reprogramming of H3K9me3 at hominoid-specific retrotransposons during human preimplantation development. Cell stem cell 29, 1031–1050.e1012 (2022).

81. P. Turelli et al., Primate-restricted KRAB zinc finger proteins and target retrotransposons control gene expression in human neurons. Sci Adv 6, eaba3200 (2020).

82. P. A. Johansson et al., A cis-acting structural variation at the ZNF558 locus controls a gene regulatory network in human brain development. Cell stem cell 29, 52–69 e58 (2022).

83. S. Yokobayashi et al., Inherent genomic properties underlie the epigenomic heterogeneity of human induced pluripotent stem cells. Cell reports 37, 109909 (2021).

84. E. Deneault et al., Complete Disruption of Autism-Susceptibility Genes by Gene Editing Predominantly Reduces Functional Connectivity of Isogenic Human Neurons. Stem Cell Reports 11, 1211–1225 (2018).

85. J. Klaus et al., Altered neuronal migratory trajectories in human cerebral organoids derived from individuals with neuronal heterotopia. Nature medicine 25, 561–568 (2019).

86. T. Sawada et al., Developmental excitation-inhibition imbalance underlying psychoses revealed by single-cell analyses of discordant twins-derived cerebral organoids. Molecular psychiatry 25, 2695–2711 (2020).

87. C. A. Trujillo et al., Reintroduction of the archaic variant of NOVA1 in cortical organoids alters neurodevelopment. Science 371 (2021).

88. A. A. Pollen et al., Establishing Cerebral Organoids as Models of Human-Specific Brain Evolution. Cell 176, 743–756 e717 (2019).

89. F. Mora-Bermudez et al., Differences and similarities between human and chimpanzee neural progenitors during cerebral cortex development. eLife 5, e18683 (2016).

90. A. E. Trevino et al., Chromatin accessibility dynamics in a model of human forebrain development. Science 367, eaay1645 (2020).

91. A. Gordon et al., Long-term maturation of human cortical organoids matches key early postnatal transitions. Nature neuroscience 24, 331–342 (2021).

92. A. Yanagida et al., Naive stem cell blastocyst model captures human embryo lineage segregation. Cell stem cell 10.1016/j.stem.2021.04.031 (2021).

93. S. Tarazi et al., Post-Gastrulation Synthetic Embryos Generated Ex Utero from Mouse Naïve ESCs. Cell 10.1016/j.cell.2022.07.028 (2022).

94. Y. Fan et al., Generation of human blastocyst-like structures from pluripotent stem cells. Cell Discov 7, 81 (2021).

95. G. Amadei et al., Synthetic embryos complete gastrulation to neurulation and organogenesis. Nature 10.1038/s41586-022-05246-3 (2022).

96. M. Yang et al., Depletion of aneuploid cells in human embryos and gastruloids. Nature cell biology 23, 314–321 (2021).

97. T. W. Kim et al., Biphasic Activation of WNT Signaling Facilitates the Derivation of Midbrain Dopamine Neurons from hESCs for Translational Use. Cell stem cell 28, 343–355 e345 (2021).

98. J. Piao et al., Preclinical Efficacy and Safety of a Human Embryonic Stem Cell-Derived Midbrain Dopamine Progenitor Product, MSK-DA01. Cell stem cell 28, 217–229 e217 (2021).

99. M. M. Chan et al., Molecular recording of mammalian embryogenesis. Nature 570, 77–82 (2019).

100. B. Spanjaard et al., Simultaneous lineage tracing and cell-type identification using CRISPR-Cas9-induced genetic scars. Nature biotechnology 36, 469–473 (2018).

101. R. Kalhor et al., Developmental barcoding of whole mouse via homing CRISPR. Science 361, eaat9804 (2018).

102. Z. He et al., Lineage recording in human cerebral organoids. Nature methods 10.1038/s41592-021-01344-8 (2021).

103. R. E. Rodin et al., The landscape of somatic mutation in cerebral cortex of autistic and neurotypical individuals revealed by ultra-deep whole-genome sequencing. Nature neuroscience 10.1038/s41593-020-00765-6 (2021).

104. M. W. Breuss et al., Somatic mosaicism reveals clonal distributions of neocortical development. Nature 604, 689–696 (2022).

105. L. Fasching et al., Early developmental asymmetries in cell lineage trees in living individuals. bioRxiv 10.1101/2020.08.24.265751, 2020.2008.2024.265751 (2020).

106. M. Watanabe et al., TGFbeta superfamily signaling regulates the state of human stem cell pluripotency and capacity to create well-structured telencephalic organoids. Stem Cell Reports 10.1016/j.stemcr.2022.08.013 (2022).

107. T. Muthusamy, O. Mukherjee, R. Menon, P. B. Megha, M. M. Panicker, A method to identify and isolate pluripotent human stem cells and mouse epiblast stem cells using lipid body-associated retinyl ester fluorescence. Stem Cell Reports 3, 169–184 (2014).

108. E. J. Fertig et al., Gene expression signatures modulated by epidermal growth factor receptor activation and their relationship to cetuximab resistance in head and neck squamous cell carcinoma. BMC genomics 13 (2012).

109. E. J. Fertig, G. Stein-O’Brien, A. Jaffe, C. Colantuoni, Pattern identification in time-course gene expression data with the CoGAPS matrix factorization. Methods in molecular biology 1101, 87–112 (2014).

110. S. Kumar, G. Stecher, M. Suleski, S. B. Hedges, TimeTree: A Resource for Timelines, Timetrees, and Divergence Times. Mol Biol Evol 34, 1812–1819 (2017).

111. R. Neme, D. Tautz, Fast turnover of genome transcription across evolutionary time exposes entire non-coding DNA to de novo gene emergence. eLife 5, e09977 (2016).

112. N. Vakirlis, A. R. Carvunis, A. McLysaght, Synteny-based analyses indicate that sequence divergence is not the main source of orphan genes. eLife 9 (2020).

